# Super-resolution Imaging of Synaptic and Extra-synaptic Pools of AMPA Receptors with Different-sized Fluorescent Probes

**DOI:** 10.1101/096966

**Authors:** Sang Hak Lee, Chaoyi Jin, En Cai, Pinghua Ge, Yuji Ishitsuka, Kai Wen Teng, Andre A. de Thomaz, Duncan Nall, Murat Baday, Okunola Jeyifous, Daniel Demonte, Christopher M. Dundas, Sheldon Park, William N. Green, Paul R. Selvin

**Affiliations:** Department of Physics, Center for Biophysics, and Quantitative Biology, and Center for the Physics of Living Cells, University of Illinois, Urbana-Champaign; Department of Neurobiology, University of Chicago and the Marine Biological Laboratory; Department of Chemical and Biological Engineering, SUNY, Buffalo.

## Abstract

Whether AMPA receptors (AMPARs) enter into neuronal synapses, by exocytosis from an internal pool, or by diffusion from an external membrane-bound pool, is hotly contested. 3D super-resolution fluorescent nanoscopy to measure the dynamics and placement of AMPAR is a powerful method for addressing this issue. However, probe size and accessibility to tightly packed spaces can be limiting. We have therefore labeled AMPARs with differently sized fluorophores: small organic fluorescent dyes (~ 4 nm), small quantum dots (sQD, ~10 nm in diameter), or big (commercial) quantum dots (bQD, ~ 20 nm in diameter). We then compared their diffusion rate, trajectories, and placement with respect to a postsynaptic density (PSD) protein, Homer 1c. Labeled with the small probes of sQDs or organic fluorophores, we find that AMPARs are located largely within PSDs (~73-93%), and generally reside in “nanodomains” with constrained diffusion. In contrast, when labeled with bQDs, only 5-10% of AMPARs are within PSDs. The results can be explained by relatively free access, or lack thereof, to synaptic clefts of the AMPARs when labeled with small or big probes, respectively. This implies that AMPARs primarily enter PSDs soon after their exocytosis and not from a large diffusive pool of extrasynaptic AMPARs.

## Introduction

A major goal of molecular and cellular neuroscience is to describe in detail the structures that mediate synaptic function and how they change during synaptic plasticity, events that are thought to underlie learning and memory formation, as well as neurodegenerative diseases (Huganir and Nicoll, 2013; Kneussel and Hausrat, 2016; Volk et al., 2015). The position and dynamics of AMPARs and NMDA-type glutamate receptors (NMDARs), two ionotropic glutamate receptors (iGluRs), are critical for synaptic strength, as well as the changes that occur with activity. However, determining whether iGluRs in the plasma membrane are synaptic, or extrasynaptic, generally defined to be inside, or outside, respectively, of the post-synaptic density (PSD), the area capable of receiving glutamate released from the pre-synapse, has been difficult. In large part this is because the distances involved are nanometer in scale, far below the diffraction limit of visible microscopy.

One controversy has centered on the location of receptors before activation and how these receptors move into the synaptic PSDs during potentiation. For example, identifying the reservoir of receptors is a key factor to explain the mechanism behind the recovery of iGluRs from receptor desensitization and enrichment during potentiation in neurons (Kneussel and Hausrat, 2016). For this reason, the population of mobile receptors recruited during recovery and enrichment has been the subject of controversy over the last twenty years.

One possible scenario is that there is a large pool of extra-synaptic receptors on the dendrite surface membrane, which can then rapidly diffuse into PSDs (Giannone et al., 2010; Groc et al., 2007; Heine et al., 2008; Nair et al., 2013). These extra-synaptic receptors enter the synaptic cleft during potentiation by lateral diffusion at the time that the synapses are activated. Another possible scenario is that the extra-synaptic receptor pool is quite small so that what is limiting is the arrival rate of receptors from endosomal stores (Malinow and Malenka, 2002; Park et al., 2004; Z. Wang et al., 2008). Instead of the extra-synaptic pool being the primary source of new PSD receptors, it is the recycling pool of receptors in endosomes that is the main reservoir for synaptic potentiation.

In the last decade, a number of workers have pioneered work on glutamate receptor dynamics using advanced fluorescence microscopic techniques (Groc et al., 2007; Heine et al., 2008; Kneussel and Hausrat, 2016; MacGillavry et al., 2013; Nair et al., 2013). Dahan et al. found that commercial quantum dots could get into the synaptic cleft, at least occasionally, as visualized by electron microscopy (Dahan et al., 2003). (Due to the large size of commercial quantum dots, we call them big quantum dots, or bQDs.) Because the bQDs are extremely bright, the position can be localized to within ~10 nm on the sub-second time-scale via high-resolution visible fluorescence. They and other workers have since claimed that ~50% of AMPA receptors are in extra-synaptic regions on the surface of dendrite membranes and are relatively free to move via diffusion to rapidly repopulate the synapse (Heine et al., 2008; Tao-Cheng et al., 2011).

However, the bQDs are approximately the same size as the cleft, >20 nm in diameter, and the cleft is further populated by extrasynaptic matrix material. This would make it difficult for unfettered access of the bQD-iGluRs. To test if this was the case with bQD-AMPAR, in particular, we made small quantum dots (sQDs) that are below 10 nm in diameter and attached them to AMPAR via small linker proteins— either streptavidin (~ 5 nm or 58 kD) or single-chain antibody (~4 nm or 14kD) (Cai et al., 2014; Y. Wang et al., 2014). We found that the 85-93% of the AMPARs that were labeled with sQDs were highly constrained within the synaptic cleft, with the remaining being relatively free to diffuse in the extra-synaptic regions. In contrast, only about ~15% of AMPARs labeled with bQDs were constrained in the synapse, and ~85% were *free* to diffuse extra-synaptically. Presumably the inability of the bQDs to enter PSDs is because the large size of the bQDs, as opposed to the smaller sQDs, prevents the labeled AMPAR from entering the synapse. Nevertheless, there were a number of technical problems, which led some to question these results. For example, chromatic aberrations and microscope drift were not adjusted for; more significantly, yet smaller probes (e.g. organic probes) with the modern super-resolution techniques (Betzig et al., 2006; Huang et al., 2008) were not used for imaging receptors in live neurons.

Others have used single particle tracking (spt) of small (< 5 nm) fluorescent probes permanently or temporarily bound to an AMPAR. One such technique is called sptPALM, where iGluRs that were permanently bound to fluorescent proteins (FPs) were used; another technique is called uPAINT, which measures a fluorescent probe that is temporarily bound to the receptor. However, contradictory results were found. uPAINT found that AMPAR were either diffusive (Giannone et al., 2010), or stationary (Nair et al., 2013): sptPALM found that AMPAR were very diffusive. A possible explanation for these results is that sptPALM-FPs reports internally (and externally) labeled AMPAR that are free to diffuse, but that uPAINT only observed the plasma-membrane-bound AMPAR and that these are largely immobile. Similar concerns apply to FRAP data, which is sensitive to internalized AMPAR receptors (Heine et al., 2008). Thus, the population of recycling AMPARs in the membrane may have been significantly overestimated in these studies.

Given this continued controversy, we have measured parameters associated with AMPAR with different-sized fluorescent probes, ranging from the bQDs, to sQDs with different attached ligands for binding, to organic fluorophores (Figure 1). The parameters measured are the: 1) diffusion constant, 2) trajectory, 3) distance from the center of the PSD 4) changes that occurred over time after transfection and 5) upon removal of the extracellular matrix (summarized in Table 1). The conclusion is that with small quantum dots or with (even smaller) organic fluorophores, the vast majority of AMPA receptors are stuck in or nearby PSDs, forced to move in constrained diffusion in a “nanodomain”, which is much smaller than the diffusional volume of the entire PSD.

**Figure 1.**
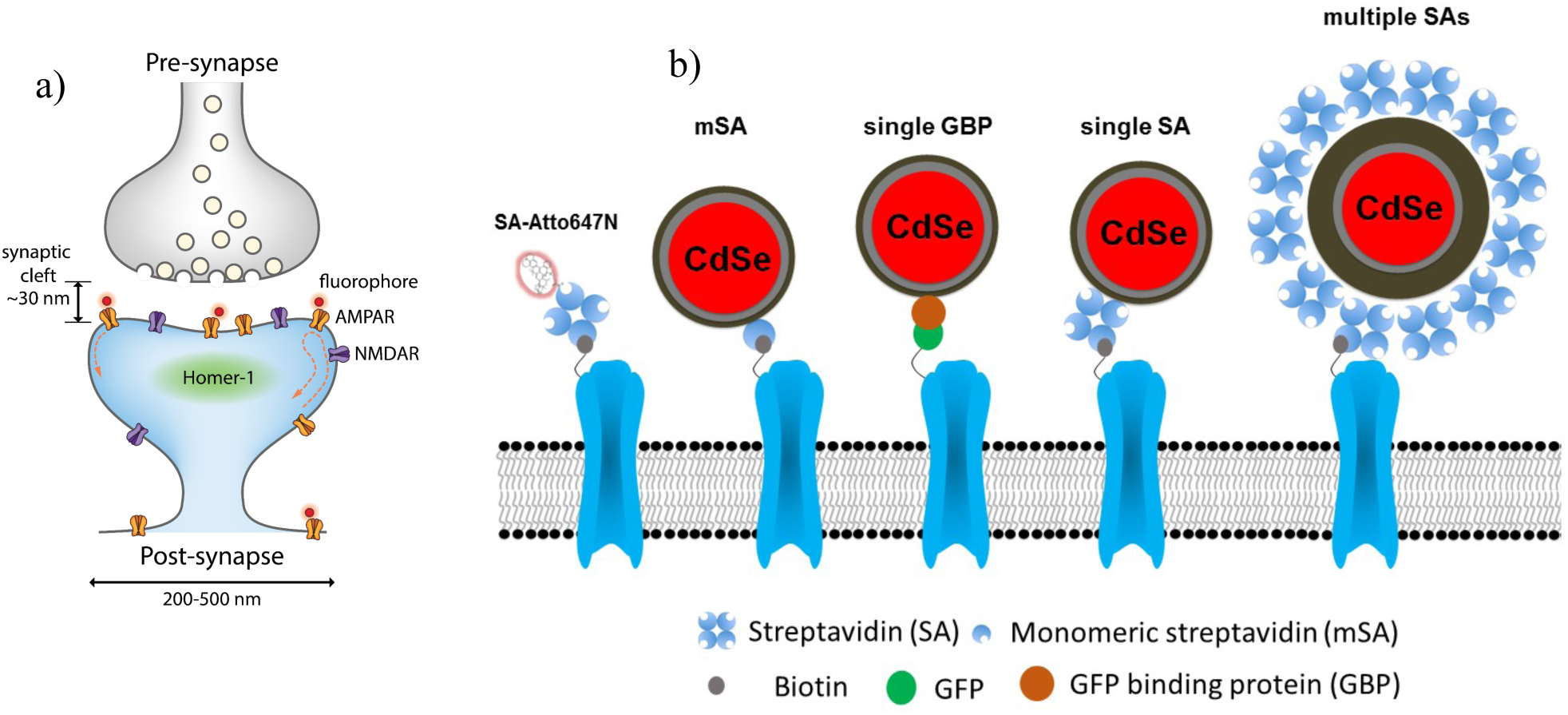
a) Schematic of synapse. Since the synaptic cleft width is only about 30 nm, small size (fluorescent) probes are required to study glutamate receptors. b) Schematics of AMPA receptors with a variety of different sized probes, with various linkers connecting them to an AMPAR. The smallest is the organic fluorophore linked by a Streptavidin (SA); then there is a small quantum dot linked by either a monomeric SA, an anti-GFP nanobody (called GBP) attached to a GFP-AMPAR, or a (single) SA; finally, there is a commercial quantum dot which comes with multiple SA.

**Table 1.** Summary of AMPA receptor diffusion measured with various fluorophores and tags.

| Fluorophore | size (nm) | tag | diffusion coefficient ($\mu\text{m}^2/\text{s}$) | trajectory range (nm) (± width) | synaptic receptors* (%) |
| --- | --- | --- | --- | --- | --- |
| Small Qdot | ~ 10 | SA | $10^{-2.41 \pm 0.02}$ | $418 \pm 172$ | 75 |
| | | GBP | $10^{-2.53 \pm 0.02}$ | $420 \pm 170$ | 82 (92) |
| | | mSA | $10^{-2.59 \pm 0.02}$ | $363 \pm 144$ | 86 (84) |
| Atto647 | ~ 5 | SA | $10^{-2.55 \pm 0.02}$ | $380 \pm 156$ | 77 (73) |
| CF633 | | | $10^{-2.70 \pm 0.01}$ | $380 \pm 140$ | 90 (93) |

\* The first and second number corresponds to the percentage of AMPAR in the lower left hand box and the area under the Gaussian curve for the trajectories, respectively, as outlined in each of Figure 3-5.

## Results and Discussion

### Labeling of AMPARs in neurons

(Figure 1) In order to label AMPARs with different fluorescent probes, different tagged versions of the AMPAR subunit GluA2 were transiently transfected into cultured rat hippocampi neurons (see Methods). These were then assayed at 13-16 days in vitro (DIV). GluA2 was genetically appended on the N-terminus with an 18 amino acid sequence, which, when cotransfected with BirA, resulted in a biotin in the N-terminus (Howarth et al., 2005). Alternatively, GluA2 with GFP on the N-terminus was also used. The AMPAR was labeled with either a small organic fluorophore attached to Streptavidin (Cy3B on fixed neurons, Figure 2c; Atto647N or CF633 on live neurons, Figure 2d); a small quantum dot with 3 different ligands (a monomeric SA, a single nanobody against GFP called GBP (Ries et al., 2012); or a SA at a ratio so that there was approximately one SA per sQD); or a big quantum dot, which came commercially with many Streptavidin molecules.

**Figure 2.**
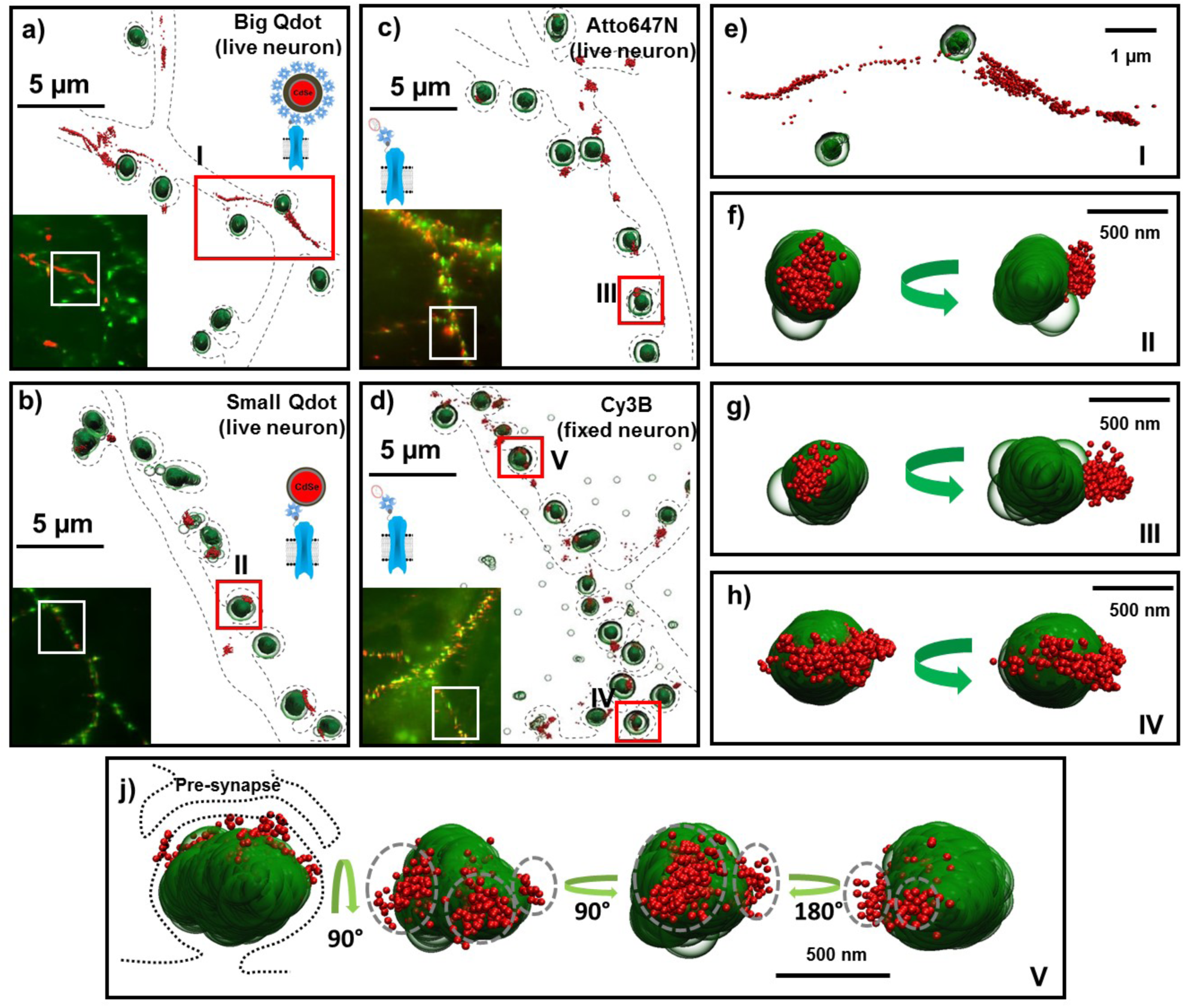
Big and Small quantum dots and organic fluorophores attached to AMPAR (red) on live cultured neurons with the PSD protein, Homer1c, labeled (green). Big (or commercial) quantum dots (a, e) bound to AMPA fail to get in the synapse and are spread extra-synaptically. Small quantum dots (b, f) and with small organic fluorophores (Atto647N) (c, g), bound to AMPAR, are stuck in nanodomains within the synapse. Fixing the neurons and using a Cy3 organic dye (d) can get the number of nanodomains per synapse; mostly there is just one (h), but occasionally there are multiple nanodomains (j). The dotted lines in a-d represent the morphology of dendrites. The green arrow (f-j) indicates the direction of synaptic cleft and the dotted line represents the shape of the synapse. Each image is the rotated structure of the synapse as represented by curved arrow. The gray dotted lines in j indicate the nanodomain of the AMPA receptors. For super-resolution images, the microscope stage drift and chromatic aberrations are corrected.

The cultured neurons were also transfected with Homer1c, a post-synaptic-density (PSD) protein with a photoactivatable fluorescent protein (mEos3.2 or mGeos). Homer1c served as a super-resolution marker for PSDs (Figure 1 and 2). On DIV 14 (called post-transfection day 1) or DIV16 (called post-transfection day 3), the AMPAR was labeled with a fluorophore (either organic dye of sQD or bQD). Labeling at DIV14 and DIV 16 allowed the concentration of AMPAR to vary, and so we could see whether the parameters defined were functions of time and presumably AMPAR concentration. For all tags, the Homer1c or AMPAR could be visualized in 3D (using a cylindrical lens) to locate the position of the synapse and to track their dynamics (Huang et al., 2008).

### Diffusion constants and trajectory range

For all tags, 2D “heat maps” was created which plots the diffusion constants versus the area that the AMPAR traveled. In general, the lower left-hand corner of the map shows slowly (or constrained) diffusion of AMPAR where the trajectory is limited; those in the upper right-hand corner show the opposite. Semi-quantitatively, we make the separate “slow” from “fast”, to be D ≤ 10^−1.75^ μm^2^/s and a trajectory range of ≤ 10^−0.1^ μm (0.79 μm), as shown in Figures 4–5. (This is somewhat arbitrary, though consistent with the literature (Constals et al., 2015) (Cai et al., 2014). Alternatively, the areas under a Gaussian curve can be chosen.) This map helps in the identification of different AMPAR diffusion rates as a function of position on the neuron membrane and as a function of time; furthermore, it helps to determine how this may change as a function of time after transfection, which, in turn, tells about possible cross-linking between different AMPARs. As mentioned previously all labels were tested after 1 day, and after 3 days following transfection (corresponding to DIV14 and DIV16, respectively). The results should *not* change—i.e. they should be time-independent— otherwise it indicates cross-linking may be occurring. Notice it does *not* tell about the correlation between the position of Homer1c and the AMPAR. This is shown separately (see, for example, Figure 2d, e, and 4f).

### Overview of results

Figure 2 shows representative traces of the results of live neurons labeled with bQD, sQD and an organic fluorophore (Atto647N), as well as a fixed neuron labeled with another organic fluorophore, Cy3B. Table 1 summarizes the results. Below we discuss in detail the results with each labeling technique. Briefly, what we find is that:

- The bQD-AMPAR are primarily free to diffuse widely outside of PSDs, and that their labeling changes with time (DIV) after the transfection of the AMPAR tagged subunits. For these reasons, the results with bQDs are difficult to quantify in terms of the distance from the center of PSDs.
- The sQD-AMPAR are labeled with 3 different ligands connecting the sQD to the AMPAR. The intent is to show ligands which do not have a tendency to cross-link, either within the tetrameric AMPA receptor, or between the AMPARs. As expected, they show no time-dependent changes after transfection. The labeled AMPAR are almost totally in, or nearby, PSDs marked by Homer1c. All sQDs in PSDs are undergoing constrained diffusion, over a small (sub-synaptic) range (+/- ~0.4 μm), generally called a “nanodomain” (Nair et al., 2013).
- Similar results to the sQDs are found with organic fluorophores. On fixed cells, we can examine the number of nanodomains per synapse. Most often, there is a single nanodomain per synapse, although occasionally there are multiple ones.
- If the extracellular matrix (ECM) is removed, the diffusion of AMPARs with sQDs does not change while the diffusion of AMPARs with bQDs does significantly change.

These results suggest that the large diffusive pool of extra-synaptic AMPARs observed when labeled with bQDs is largely an artifact caused by the inability of most bQD-tagged AMPARs to enter synaptic clefts. These results argue that AMPARs entering and exiting PSDs from the plasma originate from or reenter the intracellular pool of recycling AMPARs.

### AMPAR-bQD

Most of the AMPA receptors labeled with SA-bQD on DIV14, i.e. one day following transfection, reside in a widely spread extra-synaptic region (Figure 2a and e). This is consistent with our previous results (Cai et al., 2014) with the difference that the microscope stage drift and chromatic aberrations are corrected in this study (supplementary information and Figure 2 – figure supplement 1~7). In addition, we plotted a 2D-map, for one-day and three- days after transfection, respectively (Figure 3a, b). For the bQDs, the labeling is a function of DIV, indicating that the diffusion time and trajectory is a function of time. Presumably, this is because of cross-linking of AMPAR-bQD, which would depend on the amount of AMPAR expressed in the cell membrane. As a result, the results with bQDs are difficult to determine the diffusion or the position of AMPARs in neurons.

**Figure 3.**
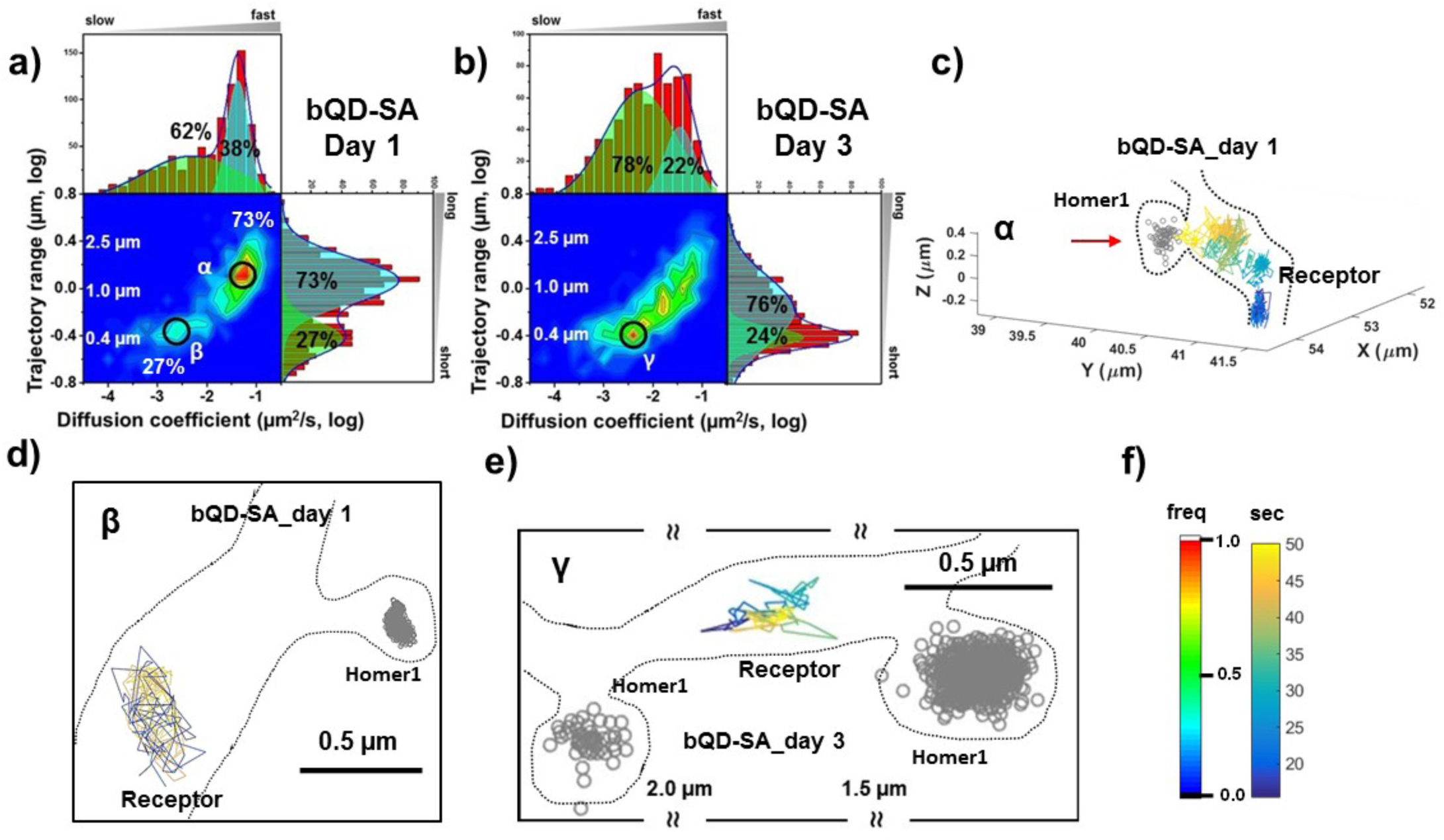
AMPARs are *not* synaptic as measured by QDs. Two-dimensional plot of the diffusion coefficient and the trajectory range of AMPA receptors using big qdots (a) measured day 1 (n = 483 traces) and (b) day 3 (n = 715 traces) after transfection. White dot lines are for 10^−1^ μm^2^/s (0.79 μm) and 10^−1.75^ μm^2^/s (0.018 μm^2^/s)) for the trajectory range and the diffusion coefficient, respectively. Right upper side represents the population of fast diffusive receptors with longer trajectory and the left lower side represents the population of slow diffusive receptors with shorter trajectory. For all histograms, the distribution could be deconvoluted by Gaussian functions. Cyan colored Gaussian functions represent the fast diffusive receptors in diffusion coefficient and longer diffusive population in trajectory range. Green shaded Gaussian functions indicate slower and shorter diffusive receptors. On Day 1, the fast and long-distance peak (α) corresponds to ~73% of all molecules with ~ 10^−1.25^ μm^2^/s (α) and ~1.3 μm, while the slow and short diffusion population (β) corresponds to ~ 27% of all molecules with ~ 10^−2.5^ μm^2^/s and ~ 0.4 μm. On day 1, the α peak is not in the synapse but diffusive relatively freely along the synapse as represented in (c). Dotted lines in c) represent the morphology of dendrite and the red arrow in c) indicates the synaptic cleft. The remaining 27 % on day 1 in figure 3a has a short trajectory and > 10 x slower diffusion than the α peak. The majority of the slow diffusive receptors on both Day 1 and Day 3 are located away from synapse as in d) and e). At day 3 after transfection, the diffusion coefficient and trajectory range measured by using big qdots are significantly changed, probably due to crosslinking. f) shows the color coding for population of heat maps and for time traces for a single receptor.

At post-transfection day 1, Figure 3a shows there are primarily two peaks with the majority (73%, peak α) having a fairly fast diffusion constant (10^−1.25^ = 0.056 μm^2^/s) with about a ~1.3 μm travel length, and a minor peak (27%, peak β) having a fairly slow diffusion (10^−2.5^ = 0.0032 μm^2^/s) with a relatively immobilized trajectory range (0.5 μm). The α peak is not in the synapse, undergoing free diffusion (in dendrites); these AMPA receptors are also not co-localized with a nearby Homer1c (Figure 3c). The β peak of AMPAR has two components, both of which are slow: 5% of AMPAR are colocalized with Homer1c, and the other 22% of AMPAR are immobilized without having a labeled-Homer1c present (Figure 3d). This 22% fraction of AMPAR may be due to Homer1c not being labeled with an FP, or an AMPAR cross-linked in extra-synaptic regions.

On post-transfection day 3, Figure 3b shows an overall broad distribution of the diffusion coefficients and trajectory range, covering the range seen at post-transfection day 1. Unlike the result at post-transfection day 1, the population of free diffusion (i.e. the α peak) has significantly decreased, while the population of slow diffusion (i.e. the β peak) has increased. The data also show that the fraction of the slow diffusion is significantly increased on post-transfection day 3, as compared with that on post-transfection day 1. A representative trajectory is shown in Figure 3e, which shows the peak γ of the bQD undergoing limited-diffusion in a small region far from synapses. We believe that this is due to the cross-linking of bQDs on post-transfection day 3, because the SA-bQD has multiple binding sites to biotin-AMPA receptors (as shown in Figure 1) and there are presumably more receptors present on day 3 than day 1. Hence, labeling with bQDs is problematic because of crosslinking to the many streptavidins, as well as size.

### sQD linked through SA, GBP, or mSA

We labeled AMPARs with sQDs that were approximately half the diameter of the bQDs or 1/8^th^ the volume. To enable site-specific attachment to the Biotin-AMPAR or to GFP-AMPAR, the sQDs were conjugated with one of three linkers for attachment to the AMPARs. They were: 1) a limiting amount of SA such that each sQD has approximately one SA; 2) a 13kD single-chain antibody against GFP (from alpaca, commercially known as the Green Fluorescent Binding Protein, or GBP), which has only one target on a GFP_N-term_-AMPAR (Ries et al., 2012; Rothbauer et al., 2008; Y. Wang et al., 2014). (It has also been directed to a pHluorin-AMPAR, a GFP-derivative (Ashby, 2004). The result for this construct is the same as GFP_N-term_-AMPAR); 3) a monomeric Streptavidin, which has only one binding site for biotin and was further molecular engineered from a recent one (Chamma et al., 2016; Lim et al., 2012; 2011), in order to minimize non-specific labeling with neurons (Figure 4 –figure supplement 1). The SA and mSA were used with the same AMPAR constructs as used with the bQDs.

#### Slow motion of AMPAR-sQDs

The results for a 2D “heat map” show that the results from post-transfection day 1 and from post-transfection day 3 within a particular linker-type are essentially identical for each of the sQDs attachment methods (Figure 4a-c; summarized in Table 1). This implies that cross-linking of the sQDs to AMPAR is not a problem. Furthermore, all three data sets predominantly show slow diffusion, with a diffusion coefficient ~ 10^−2.5^ μm^2^/s. (10^−2.41±0.02^ μm^2^/s for SA; Figure 4a; 10^−2.53±0.02^ μm^2^/s for GBP, Figure 4b; 10^−2.59 ± 0.02^ μm^2^/s for mSA, Figure 4c). This yields 75% (70%), 82% (92%) and 84% (86%) slowly moving, i.e. *a vast majority display constrained diffusion.* (The numbers in parenthesis correspond to taking the integrated area under the Gaussian fit). By looking at the diffusion constants, one sees a distribution of constrained diffusion: 84%, 100%, 96%. Both results are in direct contrast with the results for labeling with the bQDs. In addition, the diffusion constant for AMPAR linked via SA to sQD is somewhat broader than the sQD coupled to GFP via GBP, which is somewhat broader than the mSA. (The mSA result is approximately equal to than the organic fluorophores—see below). This possibly implies that, although SA-sQDs are significantly better than the bQD to label AMPARs, smaller ligands (like mSA) or organic-fluorophores with SA (see below) are better to study synaptic receptors in neurons.

**Figure 4.**
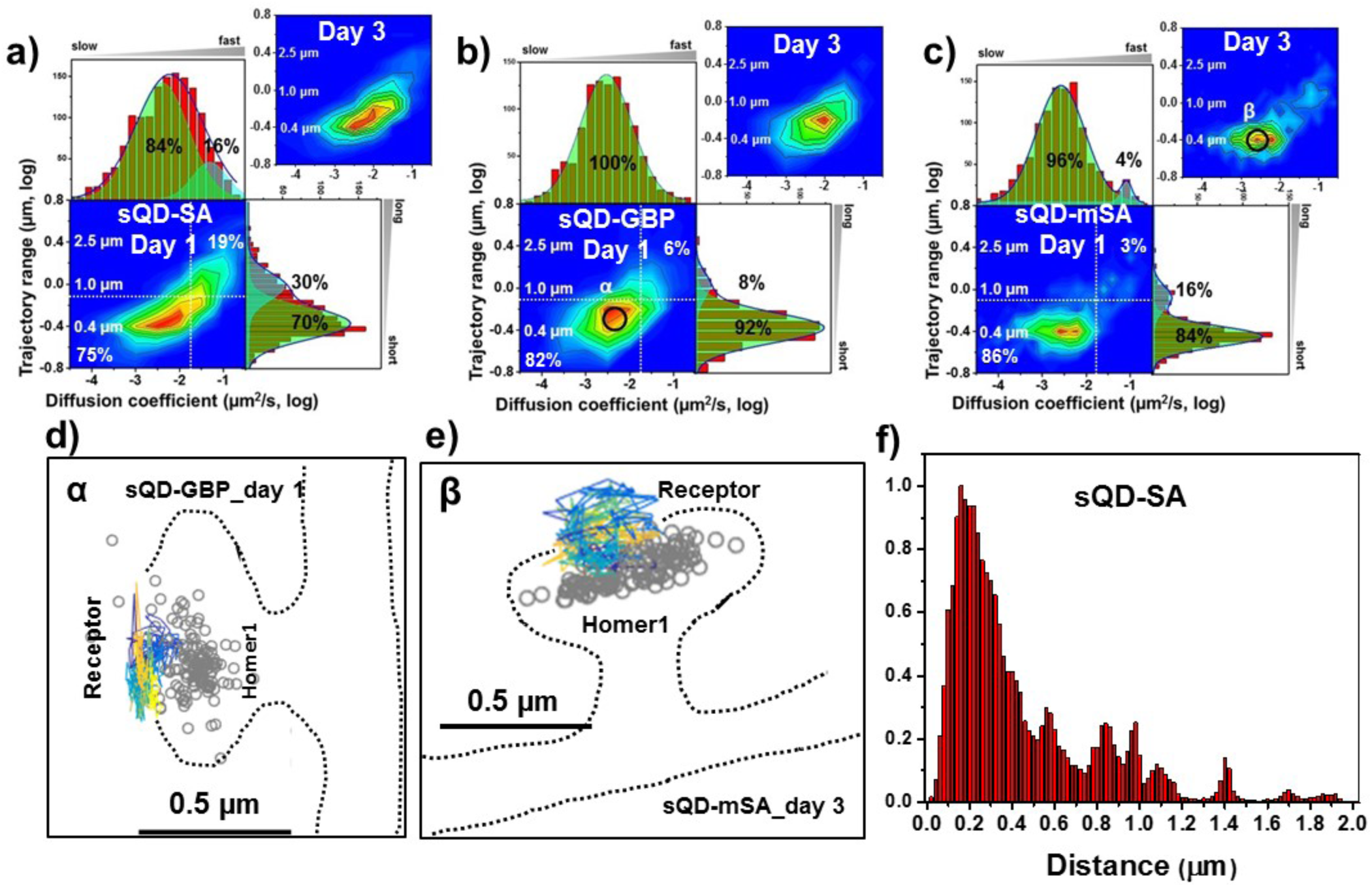
AMPARs are synaptic as measured by sQDs. Two-dimensional plot of the diffusion coefficient and the trajectory range of AMPA receptors using small qdots with various tags, (a) SA (n = 1453 and 1523 traces for day 1 and 3, respectively), (b) GBP (attached to a GFP-AMPAR) (n = 524 and 537 traces for day 1 and 3, respectively), and (c) mSA (n = 750 and 701 traces for day 1 and 3, respectively), measured day 1 and day 3 after transfection. White dot lines are for 10^−1^ μm^2^/s (0.79 μm) and 10^−1.75^ μm^2^/s (0.018 μm^2^/s)) for the trajectory range and the diffusion coefficient, respectively. Upper right side represents the population of fast diffusive receptors with longer trajectory and lower left size represents the population of slow diffusive receptors with shorter trajectory. Fitting the graphs via two Gaussian functions, the cyan represent the fast diffusive receptors (~ 10^−1.25^ μm^2^/s) in diffusion coefficient and longer diffusive population (1 μm) in trajectory range; green indicates slower (~ 10^−2.5^ μm^2^/s) and shorter diffusive receptors (~ 400 nm). For all three tags, 75% - 86% are slowly diffusive and have a short trajectory. Insets in a), b), and c) are two-dimensional plot for diffusion coefficient and trajectory range, measured day 3 after transfection. All insets are similar to the result, measured day 1. Slow diffusive receptors are mainly localized in synapses as in d) (peak α in b) and e) (peak β in c). f) histogram for the distance between receptors and the center of Homer1c. The histogram shows that 75% of receptors are localized within 0.5 μm. Color-coding for heat maps and for time traces for a receptor is the same as Figure 3f.

#### Distance of AMPAR to Homer1c

We also measured the distance of the AMPAR to the center of Homer1c. Qualitatively, the receptors are often in “nanodomains,” sub-synaptic regions where the receptor can diffuse slowly (Figure 2 b, f; also Figure 4d, 4e). This has been previously seen using sptPALM and uPaint (Nair et al., 2013), although our results on sQDs, because of the tremendous photostability of sQDs, are for continuous measurements over 50 seconds. As shown in Figure 4f, the distance between receptors and the center of Homer1c has a maximum at 150 ± 42 nm, with 75% of receptors being localized within 0.5 μm of Homer1c. This implies that the slow motion is in fact, slow motion of an AMPAR-sQD around, or inside of, the PSD

We also need to take into account those sQDs that are bound to AMPAR but not in a synapse. (The figure was 22-5% = 17% for the bQDs.) We find that there is 12% of the mSA-AMPAR that is constrained diffusion distant from labeled Homer1c PSDs. Again, whether this is due to AMPAR constrained within PSD not labeled by Homer1c, or constrained within some other domain, is not known.

### AMPAR in live neurons labeled with organic dyes

Here, we employed organic dyes to observe the diffusion of AMPA receptors in live neurons (Figure 2c, 2g; summarized in Table 1). This allows a direct comparison of AMPA receptors labeled with bQD and sQD and those labeled with organic fluorophores in live neurons. By choosing fairly photostable dyes, uPAINT on membrane-bound receptors can be done, thereby avoiding the problems with internally labeled receptors in sptPALM. It also allows a relatively long time (limited by the photobleaching time) for the motion to be measured.

We labeled AMPAR using SA-Atto647N or SA-CF633 (Figure 5). Both of these dyes are bright, especially when several dyes are conjugated per streptavidin, and fairly photostable (> 50 sec observation time) (Figure5 –figure supplement 1). In addition, Atto647N has a positive charge and is relatively hydrophobic, whereas CF633 is negatively charged and hydrophilic (Zanetti-Domingues et al., 2013). Somewhat more non-specific labeling was shown with Atto647N. Overall, these factors have been found to be significant in terms of the measurement of the diffusion of membrane proteins (Zanetti-Domingues et al., 2013).

**Figure 5.**
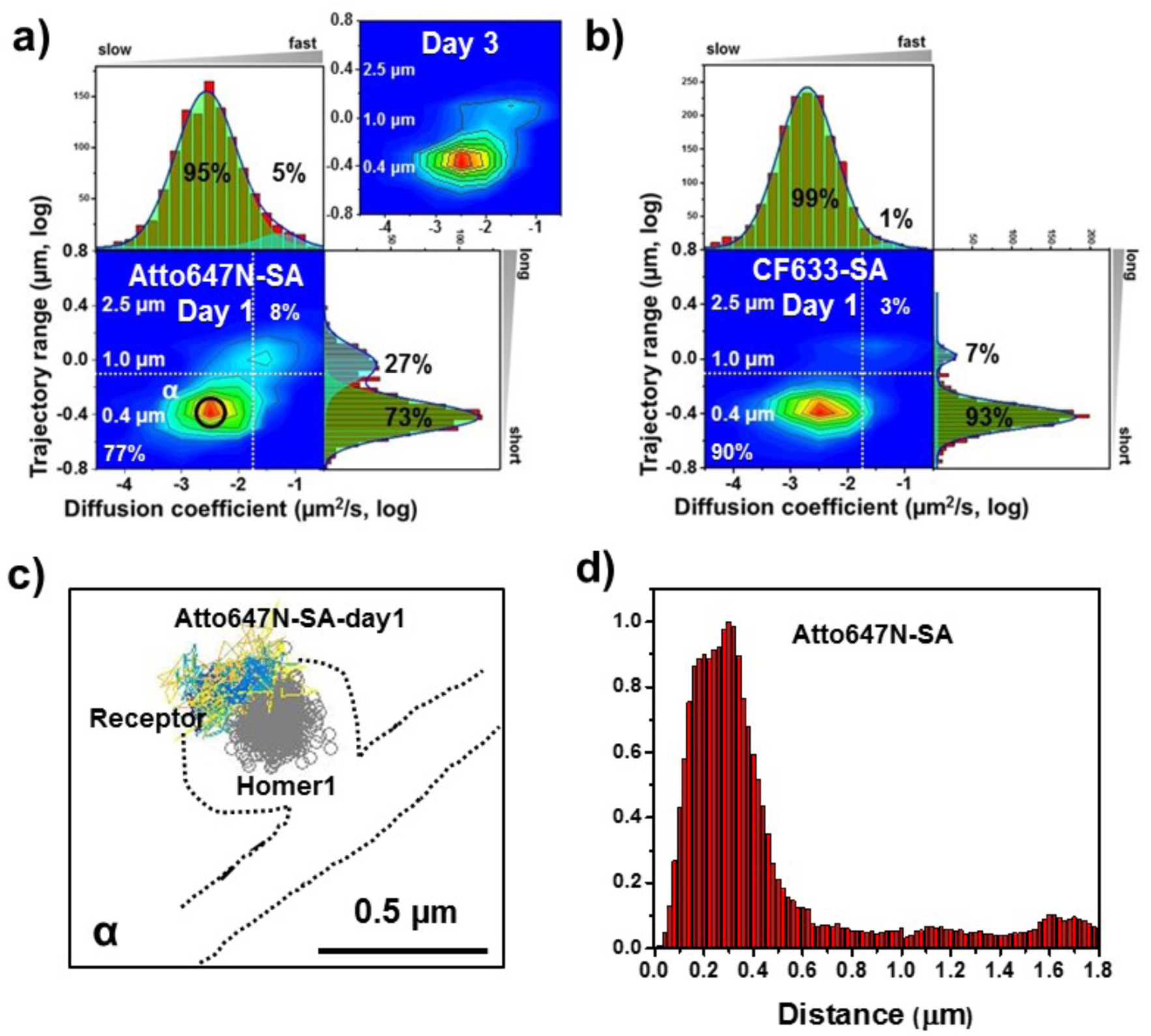
AMPARs are synaptic as measured by organic fluorophores. Two-dimensional plot of the diffusion coefficient and the trajectory range of AMPA receptors using 1 nM organic dyes for labeling, (a) Atto647N measured at day 1 as well as day 3 (n = 1152 and 3063 traces for day 1 and 3, respectively) and (b) CF633 measured at day 1 (n = 1599 traces). White dot lines are for 10^−1^ μm^2^/s (0.79 μm) and 10^−1.75^ μm^2^/s (0.018 μm^2^/s)) for the trajectory range and the diffusion coefficient, respectively. Upper right side represents the population of fast diffusive receptors with longer trajectory and lower left size represents the population of slow diffusive receptors with shorter trajectory. Cyan colored Gaussian functions represent the fast diffusive receptors (~ 10^−1.25^ μm^2^/s) in diffusion coefficient and longer diffusive population (1 μm) in trajectory range. Green shaded Gaussian functions indicate slower (~ 10^−2.5^ μm^2^/s) and shorter diffusive receptors (~ 400 nm). For both organic dyes, more than 90% have a small diffusion constant and have a 73-93% have a short trajectory (based on the diffusion coefficient graph). c) We find that they are moving slowly next to the Homer1c. d) histogram for the distance between receptors and the center of Homer1c on a live neuron. The histogram shows that 78% of receptors are localized within 0.5 μm. The color-coding for heat maps and for time traces for a receptor is the same as Figure 3f.

#### Constrained diffusion in nanodomains when AMPAR is labeled with organic fluorophores

Figure 2c shows an example of AMPARs labeled with organic-fluorophores on the live cells are colocalized with Homer1c in nanodomains. A 2D “heat” map (Figure 5a and 5b) shows that: the diffusion constants and range of AMPAR are both quite small when labeled with both Atto647N (post-transfection day 1 and day 3) and CF633 (post-transfection day 1). The results for Atto647N at various concentrations (0.1-1 nM) are similar (Figure 5 –figure supplement 2). This indicates that AMPAR is limited to constrained diffusion in nanodomains, With Atto647N, there is approximately a quarter of the AMPAR which has longer trajectory range and larger diffusion constants; these are not much present with the CF633 dye, where we find that 93% are constrained in nanodomains. Finally, figure 5c and 5d indicates that the AMPAR is adjacent to the range of Homer1c with 78% of receptors localized with 0.5 μm to Homer1c.

#### AMPAR in fixed neurons with organic dyes

To measure the overall population of AMPAR within one synapse, as opposed to the behavior of an individual AMPAR, we obtained super-resolution (direct-STORM) images for neuronal receptors using a heavily labeled organic fluorophore (SA-Cy3B) after fixing the neurons (Figure 2d, 2e) (Heilemann et al., 2008). Cy3B is particularly good with STORM because it can be made to photoactivatable with NaBH_4_ (Vaughan et al., 2012). Furthermore, we can label extensively with many organic fluorophores because the hydrodynamic diameter of organic fluorophores is ~ 1 nanometer and even when conjugated with streptavidin, they are only about 4 nm in diameter (Dikić et al., 2012). Consequently, SA-organic dyes are significantly smaller than either bQDs or sQDs or even labeled-antibodies (which are commonly used to label AMPAR). In addition, non-specific labeling was insignificant (Vaughan et al., 2012).

Most receptors are localized at about 130 ± 61 nm away from the center of Homer1c (Figure5 –figure supplement 3). Furthermore, 85% of the receptors are localized within 0.5 μm of Homer1c, indicating the close association between the Homer1c and AMPARs. These findings mirrored those obtained using sQDs and organic dyes on live neurons, namely that the localization of AMPA receptors is predominantly at the synaptic regions. In addition, we observe in these fixed samples, that there is typically one nanodomain per synapse (Figure 2h), although multiple nanodomains do exist (Figure 2j).

#### Removing extra-cellular matrix

The extra-cellular matrix (ECM) is a dense meshwork structure in surrounding brain cells (Syková and Nicholson, 2008). In neurons, surface receptors such as AMPA and NMDA receptors, possibly interact directly or indirectly with the ECM, so that the ECM may affect the lateral diffusion of surface receptors in the synapses and surrounding locations (Dityatev and Schachner, 2003). Recent studies have shown that the ECM is also involved in the mechanism of recruiting glutamate receptors through lateral diffusion (Dityatev and Schachner, 2003; Dityatev et al., 2010; Frischknecht et al., 2009) (Frischknecht and Gundelfinger, 2012). In particular, one recent study claimed that when the ECM was enzymatically removed, the extra-synaptic AMPA receptors moved at a higher diffusion rate and the exchange of synaptic receptors was increased (Frischknecht et al., 2009). They therefore concluded that the ECM might regulate short-term synaptic plasticity. However, they used bQDs, so their conclusion needs to be re-examined.

We therefore compared a normal neuron with ECM with a neuron in which ECM was removed (see Methods). This was done with both bQD and for sQD on AMPAR. Recall that with bQD, in the presence of ECM (Figure 3a and dotted lines in Figure 6a), we have two populations of AMPAR-bQD: 73% are fast moving (D ~ 10^−1.20^ ^± 0.01^ = 0.063 μm^2^/s) over a large distance and 27% are slow moving (D ~ 10^−2.0^ - 10^−3.0^ μm^2^/s), and constrained to be approximately 0.4 μm. After ECM removal, we find that the slow moving ones (the β peak in Fig. 3a or the white dotted line in Figure 6b) is gone. All of the remaining AMPAR-bQDs move fast and far. Evidently, the AMPA receptors cross-linked far from the synapses (Figure 3d), was gone without the ECM.

In contrast, with small qdots, we observed that the average and distribution of the diffusion coefficient of AMPA receptors was almost the same before (D_ave_ = 0.015 ± 0.0006 μm^2^/s) (Figure 6b, dotted line) and after (D_ave_ = 0.013 ± 0.0004 μm^2^/s) ECM removal (Figure 6c). The average rates are shown in Figure 6c. The conclusion is that the synaptic AMPARs are not affected by removal of ECM, presumably because their diffusion is limited through their interaction with PSD scaffold proteins.

**Figure 6.**
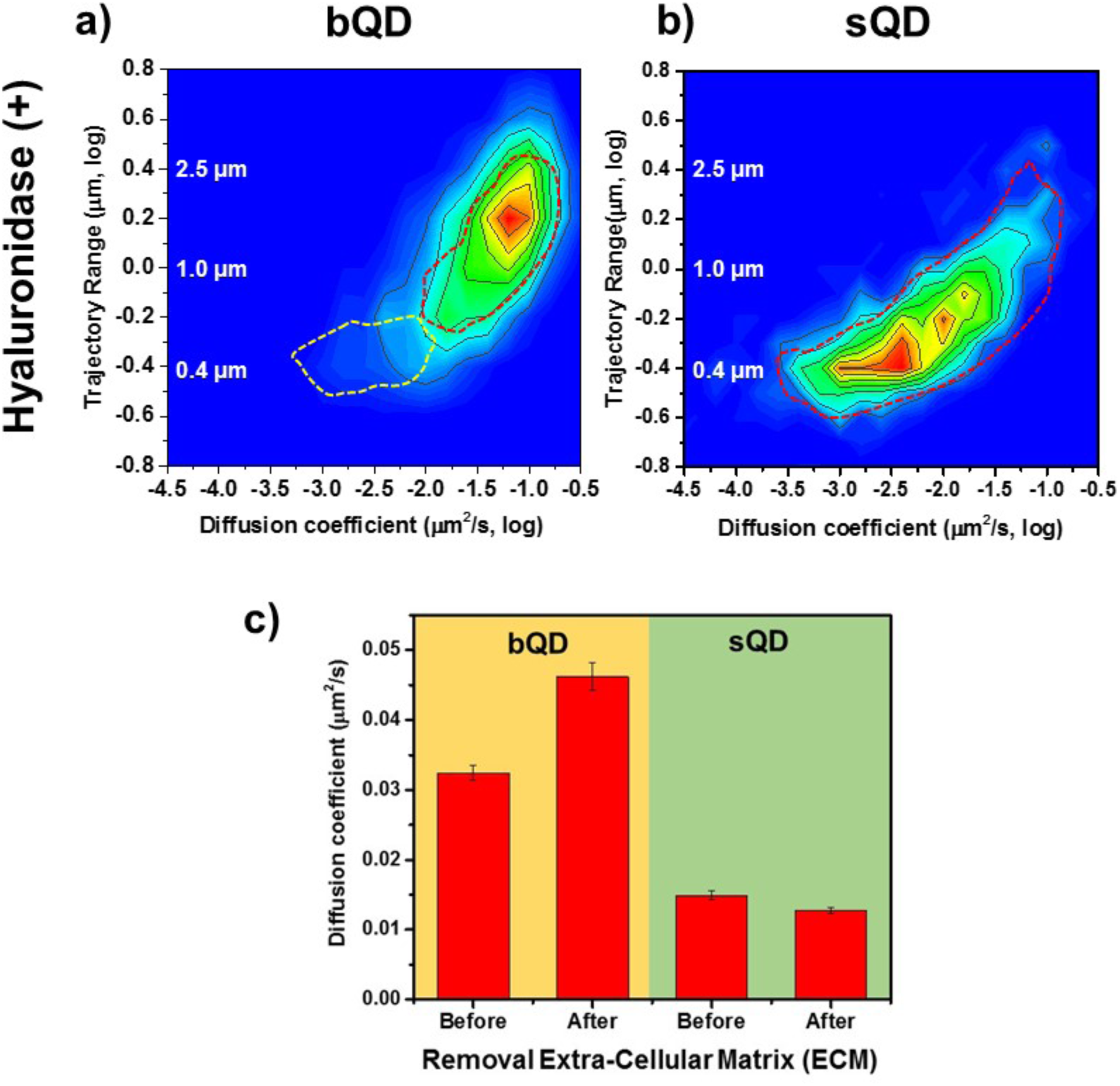
Lateral Diffusion of Synaptic AMPAR is Unaffected by Removal of ECM. Measurement the diffusion coefficient and trajectory range of the AMPA receptors after enzymatically removing extra-cellular matrix (ECM) using Hyaluronidase (100 unit/ml). (a) and (b) are 2D plots for diffusion coefficient and trajectory range measured with big qdots (n = 2566 traces) and small qdots (n = 1103 traces), respectively, after removal ECM. Dot lines circles represent the diffusion coefficient and trajectory range of Hyaluronidase (-) (taken from Figure 3a and Figure 4a, respectively). When using big qdots, the population of slow diffusion and short trajectory range disappears with treating with Hyaluronidase. However, with sqdots, there is no significant difference. The color coding for heat maps is the same as Figure 3f. c) The average diffusion coefficient, measured by using big qdots, after the removal of the ECM increases 1.4 fold from that at the condition of Hyaluronidase(-). On the contrary, the average diffusion coefficient at Hyaluronidase (+) is almost the same as without treating Hyaluronidase.

#### Agreement with previous studies

In contrast to the studies showing that AMPAR is highly diffusive along the dendrites, several studies, using various techniques, have shown that AMPA receptors are mostly localized at synapses. First, electrophysiology allows us to estimate the population of receptors in the synapses as well as in the dendrites. Cottrell *et al.* observed the density of AMPARs by measuring the glutamate-evoked response current through glutamate stimulating in synaptic regions or in dendrites. They observed the high current signal after glutamate stimulation in synaptic regions, but very low current signal with stimulation in extra-synaptic regions. Through comparing signal strength, they concluded that less than 1% of AMPA receptors were populated in the extra-synaptic regions after synaptogenesis by 11 days (or more) *in vitro* (DIV) (Cottrell et al., 2000). Second, studies using fluorescence microscopy showed that AMPARs (MacGillavry et al., 2013) were co-localized with PSD-95 in dissociated cultured neurons, indicating that they were heavily colocalized with the synaptic regions. In addition, it was reported that AMPA receptors were mostly localized at the synaptic region in hippocampal neurons (Passafaro et al., 2001). Here they observed that more than 75% of AMPA receptors, both subunits GluA1 and GluA2, were colocalized with Shank at the synaptic regions. Also, their spatial and temporal fluorescence images showed that the exocytosed receptors accumulated over time (~ 30 min) at the synaptic regions. Lastly, electron microscopy (EM) is another way to directly observe the location of receptors in neurons. EM was used to show that AMPA receptors were localized at synapses (Kharazia et al., 1996). All these observations indicate that most glutamate receptors are localized and clustered in the synaptic active zone in mature neurons.

## Conclusion

In general, when synapses are excited or stimulated, more AMPA receptors are required to receive more signals. Even under basal conditions, the AMPARs are constantly moving in and out of the PSD. Consequently, it is important to identify where the pools of AMPA receptors are located in order to understand the mechanism for AMPA receptors to enter into the synaptic cleft. As we described above, there are two possibilities for where the AMPA receptors originate: one is that a significant number of receptors are located in the membrane away from the synapse (somewhere on the dendrite) and diffuse in when needed; the other is that AMPA receptors outside PSDs are largely in intracellular pools and undergo exocytosis to supply new synaptic AMPARs. In this study, we found very little evidence for a sizeable pool of extrasynaptic receptors coming in from the dendrite. Specifically, we found that 84% ~ 97% of AMPA receptors were immobilized within PSDs (which included immobilized AMPA receptors without labeled Homer1c-mGEOS, approximately 10% ~ 15% of total AMPAR), and only 5% ~ 16% were free diffusive along the dendrites. From these results we conclude that the majority of AMPA receptors outside of the PSDs do not come from cell-surface pools but rather from intracellular pools.

## Acknowledgements

This work was partially supported by NIH NS087413 to PRS and WNG, by NIH NS097610 and NSF PHY-1430124 to PRS and NSF CBET-1264051 to SP.

## Methods

### Microscope

Experiments were performed with a Nikon Ti Eclipse microscope with a Nikon APO 100 X objective (N.A. 1.49). The microscope stabilizes the sample in z-axis with the Perfect Focus System. An Agilent laser system MLC400B with 4 fiber-coupled lasers (405 nm, 488 nm, 561 nm and 640 nm) was used for illumination. Elements software from Nikon was used for data acquisition. A back illuminated EMCCD (Andor DU897) was used for recording. For 3-D imaging, a cylindrical lens (CVI Melles Griot, SCX- 25.4-5000.0- C-425-675) of 10 m focal length was inserted below the back aperture of the objective. A motorized stage from ASI with a Piezo top plate (ASI PZ-2000FT) was used for x-y-z position control. A quad-band dichroic (Chroma, ZT405-488-561-640RPC) was used and band-pass emission filter 525/50, 600/50, 710/40, 641/75 was used for fluorescence imaging.

### Controlling for Stage drift and chromatic aberration correction

Stage drift and chromatic aberration are well known and pose significant problems in super-resolution images. As we reported earlier (Lee et al., 2012), making fiduciary-marked patterns on coverslips is an efficient method to correct for stage drift. Here we modified the procedure to work with cultured neurons. Chromatic aberration, when using several different dyes that emit at differing wavelength is also a problem. Using a nanohole pattern and differing wavelengths of excitation, we have been able to reduce the error of chromatic aberration to less than 2 nm, as described (DeWitt et al., 2012; Pertsinidis et al., 2013; Y. Wang et al., 2014). Here we also find that by taking 3D images (−400 nm < z < 400 nm), the mapping function taken at z = 0 is valid within 4 nm. These techniques enable 3D multi-color super-resolution imaging with minimal errors. (SI Figure S7)

### Cell culture and labeling

Primary hippocampal cultures were prepared from E18 rats according to UIUC guidelines as previous described (Cai et al., 2014) with the following modifications. In brief, hippocampal tissues were dissociated in 3 mg/mL protease and plated on 25 mm coverslips coated with 1 mg/mL poly-l-lysine and laminin. Neurons were cultured at 37 °C with 5% CO_2_ in neurobasal media with B-27 supplement, 2 μM glutmax and 50 unit/mL penicillin and 50 unit/ml streptomycin. On 12-13 days in vitro (DIV), neurons were co-transfected with Homer1c-mGeos (1 μg/coverslip) with GluA2-AP (1 *μ*g/coverslip) and BirA-ER (1 *μ*g/coverslip), and Homer1c-mGeos (1 μg/coverslip) and GluN2B-GFP (1 μg/coverslip), for AMPA and NMDA receptors, respectively, by using Lipofectamine 2000 transfection reagent. At 24 ~ 72 hours after transfection, the coverslips were transferred to warm imaging buffer (HBSS supplemented with 10 mM Hepes, 1 mM MgCl2, 1.2 mM CaCl2 and 2 mM D-glucose) for 5 min incubation and mounted onto an imaging dish (Warner RC-40LP). In the imaging dish, neurons were incubated in imaging buffer containing QDs and casein (~400 times dilution for bQDs, and ~80 times dilution for sQDs; stock solution purchased from Vector labs, SP-5020) for 5 min at 30 °C and washed with 10 ml of imaging buffer. Finally, 3 mL of imaging buffer was added to the imaging dish that was subsequently mounted on the microscope. For ECM removal, we enzymatically removed the ECM using hyaluronidase (HX0514 – Calbiochem, 100 units/ml left overnight at 37 ° with 5% CO_2_).

### Expression and purification of mutant mSA

The mSA gene was amplified by PCR and cloned into pET32 containing the OmpA signal sequence to create pET32-OmpA-mSA. The OmpA signal sequence is used to export newly synthesized protein to the E. coli periplasm, from where the protein leaks to the cell culture. A 6x-Histidine tag to allow affinity purification of secreted protein using immobilized nickel follows the signal sequence.

To express mSA, *E. coli* strain BL21(DE3) pLysS was transformed with an appropriate expression vector and plated on Luria Bertani (LB) agar containing ampicillin. Several colonies were selected from the plate for overnight growth at 37 °C in an LB starter culture containing 200 μg/ml ampicillin. The following morning, the starter culture was diluted 100 fold into Terrific Broth (TB) containing 0.05% glucose, 0.5% glycerol, 0.2% α-lactose, 1 mM MgSO4, 900 μM biotin, and 200 *μ*g/ml ampicillin. The cells were grown at 37 °C and 300 rpm until OD_600_ reached 0.3, at which point the shaker temperature was reduced to 20 °C. Once the culture has reached OD_600_ = 0.6, IPTG was added to the final concentration of 75 μM and the shaker speed was increased to 375 rpm. After 24 hr induction at 20 °C, cells were removed by centrifugation and the culture medium containing secreted mSA was collected for purification. The medium was sonicated with a 200 W sonicator (30 sec on, 30 sec off for 4 cycles at 50% amplitude capacity). The pH of the culture medium was adjusted to 7.5 using NaOH and imidazole was added to 10 mM. The medium was then centrifuged at 12,000 rpm for 20 min to remove any precipitates. The clarified medium was passed through a column packed with Ni-NTA Superflow Agarose pre-equilibrated in PBS and 10 mM imidazole. The column was then washed with PBS and 20 mM imidazole. Finally, the bound protein was eluted with 300 mM imidazole in PBS. The eluted protein was concentrated and buffer exchanged to 100 mM glycine buffer pH 2.3 using a centrifugal filter to remove bound biotin. Finally, the sample was buffer exchanged to PBS. Yields were estimated based on A_280_ measurements and purity was assessed by SDS-PAGE with Coomassie staining.

### Track AMPA and NMDA receptors

After focusing on the sample in bright-field, the Perfect Focus System was activated to minimize the sample drift in z direction. The samples were then scanned in the GFP channel (488 excitation, 525/50 emission) to locate transfected cells. A fluorescent image of the cells was taken for reference. To track the QD labeled receptors, 488 nm or 561 nm lasers was used for excitation in the hi-low-fluorescence mode with an appropriate band-pass filter for collecting the fluorescence.

### Super-resolution imaging of post-synaptic density

After the tracking experiment, the PALM (Betzig et al., 2006) experiment was carried out on the same neurons. PALM was used for super-resolution of post-synaptic density. Post-synaptic protein Homer1c was used as the PSD marker and its C- terminus was fused to photoactivatable protein mGeos. A 100 ms 405 nm laser pulse was used to activate mGeos proteins from dark to green fluorescent state. The sample was then excited with a 488 nm read out laser and emission was collected with a 535/50 band-pass filter. The cycle was repeated for 200-300 times, collecting 4000-6000 frames. The z calibration was created following method described by Huang *et al.* (Huang et al., 2008) using fluorescent beads on a glass surface and applied to both tracking and PALM data.

### STORM imaging of Cy3B

Cy3B-STORM was done following the procedure described on Joshua et al(Vaughan et al., 2012a). We labeled neurons with Cy3B-SA (2*μ*M) following the procedure of sQD labeling. We then fixed the cells with a solution of 4% PFA and 0.1% Glutaraldehyde in HBSS for 10 min. Wash with 3 tines PBS. Treat the sample with 10 mM of NaBH4 in PBS freshly prepared for 10 min. Wash with PBS. Before imaging, we add imaging buffer consisting of 5 *μ*L of PCD, 20 *μ*L of PCA, 4 *μ*L of Trolox in 471 *μ*L of T-100 (100 mM Tris at ph 8.0). Images were acquired at 10Hz using 405 nm laser at low power to do activation and 561 nm laser to excite fluorescence.

### Chromatic aberration

As for chromatic aberration, we achieved to minimize the error of chromatic aberration to less than 2 nm. We also certified the validity of the mapping function taken at z=0 for 3D imaging. We came up with that the mapping function taken at z = 0 is valid to correct chromatic aberration within the error of 4 nm when taking 3D images, plus or minus 400 within 800 nm (details in SI).

### Data analysis

For the tracking data, centroids of the all the QDs were localized in all the frames and a map of all the places QDs visited were obtained. A Matlab code was used to recover the trajectories of the QDs. In brief, the code finds locations of QDs in time t, and searches for nearby QDs in time t + 1 as the next point on the trajectory. In the 3-D single particle tracking experiment, the maximum displacement of a QD in one time step is set to be 1 *μ*m. Trajectory range was obtained by calculating the range of the trajectory in the x-, y-, and z- direction and using the maximum of the three parameters. The diffusion coefficients from the trajectories were calculated in Matlab by fitting the first 4 points of mean-square-displacement curve. For the PALM data, positions of proteins detected in each frame are localized, cluster analysis was used for identify synapses. To determine if a trajectory was synaptic, the centers of synapses were determined, and trajectories within 2 *μ*m radius of each synapse were identified. For each of these trajectories, the average distance was calculated between the center of the nearest synapse and all points on the trajectory. Any trajectories with an average distance smaller than 0.55 *μ*m were considered synaptic.

The visualizations of post-synaptic densities and sQD labeled AMPA/NMDA receptor tracking trajectories were created using VMD 1.9.2 (Humphrey et al., 1996), a widely-used program for visualizing and analyzing macromolecular structures and molecular dynamics (MD) simulation trajectories. VMD is optimized for dealing with large data sets containing millions of particles and thousands of trajectory frames, as encountered in the current study.

## Supplemental Information

### Stage drift correction

Stage drift is a significant problem in taking super-resolution images. As we reported earlier (Lee et al., 2012), making fiduciary-marked patterns on coverslips is a very efficient method to correct stage drift instead of using the previously favored fluorescent beads and gold nano-particles as fiduciary markers to correct stage drift. The perfectly stable fiduciary-marked patterns proved far superior to correct stage drift since it is impossible to perfectly immobilize beads or nano-particles on a coverslip surface, making it especially difficult to image live cells. More importantly, randomly spreading beads or nano-particles is a critical drawback for using them as fiduciary markers because it needs optimal number of fiduciary markers in the region of interest, whereas our fiduciary markers are a uniform pattern. Thus, the fluorescent beads and gold nanoparticles that have typically been used to correct stage drift are not effective, especially when imaging live cells.

We have previously reported that mammalian cells can be stably cultured on fiduciary-marked coverslips without any other coating or treatment; however, although we had no problems taking super-resolution images of mammalian cells with this fiduciary pattern, we were not able to culture dissociated neurons on fiduciary-patterned coverslips.

In general, poly-(L/D)-lysine coating is necessary to culture dissociated neurons on glass coverslips because the cell membranes of neurons are negatively charged. Poly-lysine coating is not sufficient for neurons to adhere the fiduciary marker coverslips. We then employed laminin to coat coverslips. As shown in Figure 2 - supplement 3 (a), we first coated poly-lysine after treating coverslips in oxygen plasma and then applied an additional coating of laminin on the fiduciary-patterned coverslips. We found that dissociated neurons could be stably cultured on fiduciary-patterned coverslips after coating them with both poly-lysine and laminin, sequentially, as in Figure 2 - supplement 3 (b). In terms of connection between neurons, we could obtain comparably good neurons on the fiduciary marked coverslip as we could culture them on the poly-lysine coated coverslip. More details to coat poly-lysine and laminin on the fiduciary marked coverslips were described in the method section.

As shown in Figure 2 - supplement 3 (b), (c) and (d), once neurons have been stably cultured on fiduciary-patterned coverslips, they can then be successfully transfected. Furthermore, IR illumination, as shown in Figure 2 - supplement 3 (b), is effective to allow clear observation of uniformly patterned fiduciary markers without affecting the visibility of fluorescence color channels. Hence, these coated fiduciary patterned coverslips are very useful to exclude stage drift, allowing for the culture, transfection, and super-resolution imaging and tracking of live neurons.

Here, to demonstrate the method for taking super-resolution fluorescence images, neurons were transfected with two plasmids, one allowing expression of the post-synaptic protein homer1 with a photoactivable fluorescence protein, mGeos or mEOS3.2, and the other allowing expression of glutamate receptors, AMPARs with AP tag to biotinylate. We then labeled the glutamate receptors using qdots conjugated with either streptavidin or anti-GFP nanobodies. As shown in Figure 2 - supplement 3 (c), at 530 nm ~ 700 nm of emission in wavelength, we observed fluorescence from the proteins and receptors without any auto-fluorescence from the fiduciary pattern. We could clearly observe the fiduciary pattern in the IR channel and the live cells in the visible channels, as shown in Figure 2 - supplement 3 (d). Fiduciary markers in the IR image are tracked, fitting the fiduciary markers using either the Gaussian or the Airy function. The IR camera electronically synchronizes with the visible camera so that the traces of fiduciary markers as recorded by the IR camera accurately calculates the stage drift at the time the cells are imaged. We can then correct the super-resolution cell images for stage drift accordingly.

The fiduciary marker trace corresponds to the stage drift. Thus, through this post-processing, we can correct for stage drift. Since fiduciary markers are stationary, stage drift can be much more accurately calculated using them than when using more conventional existing methods of calculating stage drift with fluorescent beads or gold nano-particles. As shown in figure S6, using fiduciary markers, stage drift can be controlled within an error of around 1 nm for x and y and 15 nm for z.

### Chromatic aberration

Another technical difficulty is chromatic aberration. The three-dimensional multi-color super-resolution imaging technique allows us to observe the interaction between two different species/types of proteins or organelles in cells; however, the color shifts that occur in different color channels is a concern to take images can be even more pronounced and problematic in super resolution images different color channels. These color shifts will cause the images to be slightly different. Even though we observe exactly the same object at different color channels, the images will be a little mismatched in different color channels. Since the refractive index of light depends on its wavelength, this chromatic aberration causes focus on an object to be detected in a little different place in each color channel. This means that we have to correct this difference chromatic aberration for studying the interaction between different proteins or organelles when using multi-color super-resolution imaging.

Most commercially available objective lenses are designed to optically minimize chromatic aberration. Based on our measurements, this optical design for objective lenses will corrects within the margin of error of about 100 nm; however, given that the spatial resolution of super-resolution microscopy is only about 20 ~ 30 nm, a margin of the error of 100 nm is a significant error. This means that achromatic optical design is not sufficient to correct chromatic aberration, especially in super-resolution imaging.

Using our method, chromatic aberration can be corrected within a margin of error of 2 nm. We accomplish this with as reported.(Wang et al., 2014) In order to correct chromatic aberration, we made a nano-hole pattern on a silver- coated coverslip. Using a thermal evaporator, a glass coverslip is coated with silver at a thickness of about 100 nm using a thermal evaporator. The nanohole pattern is produced by fast ion beam (FIB) equipment, sputtering holes of about 100 nm in size every 1.5 μm. Holes are about 100 nm in diameter and are located every 1.5 μm.

We took images for the nanoholes on three at the different color channels. As shown in supporting figure Figure 2 - supplement 7 (a) -~ (d), the positions of the nanohole patterns are slightly different on each color channel. In Figure 2 - supplement 7 (d), the red and green spots are the nanoholes in the green channel and in one of two red channels, shown by zooming in on the image of the the overlaid two color images. The significantly different position of the two sets of nanohole patterns is a result of shows the positions of red and green spots are significantly different, which is caused by chromatic aberration.

We measured the chromatic aberration between the green (GFP) 530 channel and red (RFP) channels 600 and 705channels. As shown in Figure 2 - supplement 7 (e), the difference in the location of the nanoholes in between two the nanohole pattern on channels 530 and 600 averages is about 80 nanometers. We found a similar range for the difference between nanohole locations in any two of the three-color channels. This means that, when obtaining multi-color super-resolution images, the error of co-localizations of any two colors is expected possible to be at least approximately 80 nm.

Furthermore, we also evaluated the validity of the mapping functions for 3D imaging. We obtained the mapping function at a focal point of z = 0, focal point, with an error of 2nm, and, as shown in Figure 2 - supplement 7 (e), this mapping function can be applied is valid up to within z= ± 400 nm with an error of only 4 nm. when the error is 2 nm at z = 0. Actually, we obtained the mapping function at z = 200 nm, 400 nm, 600 nm, and 800 nm and, as represented in Figure 2 - supplement 7 (e), the error is a little smaller than when using the mapping function obtained at z = 0; however, since a 3 D super-resolution image using astigmatism is not discrete along the z axis, it is not easy to have mapping functions along the z axis. At focal points above 400nm, the error dramatically increased. We obtained the mapping function at z = 200 nm, 400 nm, 600 nm, and 800 nm, as represented in Figure 2 - supplement 7 (e). At focal points above 400nm, the error dramatically increased. Although the error is a little smaller at focal points higher on the z axis than when using the mapping function obtained at z = 0, we conclude that when getting 3D images using astigmatism, the mapping function at z = 0 is accurate enough to correct chromatic aberration. This is because a 3 D super-resolution image using astigmatism is not discrete along the z axis. Hence, it would be too time consuming to try to obtain more accurate mapping functions all along the z axis.

### Monomeric Streptavidin

All streptavidin homologs have a conserved three dimensional structure, but differences in the binding pocket residues lead to differences in biotin binding. For example, the main chain root mean squared deviation between mSA and streptavidin is 0.82 Å but there is ~ 10^5^ fold difference in their binding affinities. Although extremely high affinity is not needed for labeling live cells, rapid dissociation of bound biotin interferes with detection and measurement because it results in loss of fluorescence intensity over time. As such, improving the dissociation kinetics (i.e. slowing the off rate k_off_) of mSA is important to develop a useful imaging reagent. We have previously demonstrated that replacing S25 with histidine reduces k_off_ by 7.5 fold by blocking solvent entry into the binding pocket through steric hindrance (Demonte et al., 2013). Keeping water molecules out of the binding pocket is important because they compete with mSA-biotin hydrogen bonds and lead to biotin dissociation. S25 is located in the loop between β-strands 1 and 2 (L1,2), which is found at the subunit interface of streptavidin but is solvent exposed in mSA. Therefore, blocking solvent entry in mSA in part involves recreating the lost physical barrier near L1,2.

To improve the dissociation kinetics further, we modeled the binding pocket of mSA using high affinity dimer bradavidin II (Brad2), which binds biotin with K_d_ < 10^−10^ M (Helppolainen et al., 2008). L1,2 and the binding loop L3,4 adopt nearly identical conformations in mSA and Brad2, thus allowing key residues to be reliably modeled in mSA. The Brad2 residue corresponding to mSA S25 is R17, which makes a hydrogen bond and a salt bridge with G103 and D104, respectively, to form a solvent barrier around L1,2 as shown in Figure 4 - supplement 1 (a, b). Arginine is larger than serine and histidine and is therefore more effective in blocking solvent entry. To reconstruct the solvent barrier in mSA, we mutated S25 to R, and measured the dissociation rate of the mutant mSA using fluorescence polarization spectroscopy. The dissociation rate of mSA-S25R is slower than that of mSA or mSA-S25H.

Brad2 also contains F42 in L3,4 to create a hydrophobic lid over bound biotin and trap the molecule in the binding pocket. Mutating F42 to alanine significantly reduces the affinity of interaction, indicating that the hydrophobic lid also plays an important role in biotin binding. We therefore introduced the corresponding mutation T48F in mSA-S25R to create a hydrophobic binding pocket and further reduce the dissociation rate. The measured k_off_ of mSA-S25R/T48F (mSA-RF) is 37 fold lower compared to original mSA, i.e. t_½_ = 402 min, and ~5 fold lower than the best mutant to date, i.e. mSA-S25H (Figure S8c). The single point mutant mSA-T48F resulted in only limited improvement in biotin dissociation (t_½_ = 13 min), indicating that the two mutations function cooperatively. We also tested an alternative design of the hydrophobic lid with a tryptophan, mSA-S25R/T48W/D124T (or mSA-RWT), which had similarly slow dissociation kinetics, t_½_ = 330 min (Figure 4 - supplement 1 (c)). Therefore, construction of solvent barriers through cooperative mutations in L1,2 and L3,4 can modulate the dissociation kinetics and improve the mSA-biotin interaction.

To demonstrate the use of new engineered mSA in cell labeling, we expressed a biotin acceptor peptide (AP) fused to CFP and a transmembrane helix (TM) on the surface of HEK293. We then added purified *E. coli* biotin ligase, BirA, to the cell culture to induce site-specific biotinylation of AP. The cells were fixed and labeled with mSA-RF—EGFP for imaging by confocal microscopy. The CFP positive cells were selectively labeled, suggesting high specificity of interaction between mSA-RF and biotin (Figure 4 - supplement 1 d and e). Furthermore, the intensity of EGFP fluorescence remained high after 1 hr, indicating that the interaction is stable and mSA-RF remained stably bound during measurement.

In order to verify whether SA-qdots were in fact cross-linking, we compared the diffusion coefficient of mSA-qdots and SA-qdots on the lipid bilayer at different concentrations (0.2 % and 1.0 %) of biotinylated lipid molecules. If a qdot binds to one biotinylated lipid molecule, the resulting labeled lipid molecule will diffuse at the same rate as a free diffusion of a lipid molecule; whereas, if a qdot binds to multiple biotinylated lipid molecules, the diffusion rate will significantly slow down as shown in figure S10.

We first determined whether there is any difference in the diffusion rate of molecules labeled in three ways: 1) those labeled with normal SA-qdot conjugates formed with big qdots (one conjugate has many streptavidin and numerous binding site on SA-big qdots); 2) those labeled with normal SA-qdot conjugates formed with small qdots (one conjugate has four binding sites from the streptavidin); and 3) those labeled with mSA-qdot conjugates using monomeric streptavidin and small qdots (one conjugate has only one binding site). We hypothesized that normal streptavidin conjugates with both small and big qdots would be much more likely to bind multiple biotins than would be the case with mSA conjugates. We found this to be the case, as represented in Figure 4 - supplement 3. Figure 4 - supplement 3 shows that there was little difference in the diffusion rate of molecules labeled with normal SA-qdots, whether conjugated with small or big qdots: there were a large number of slow diffusive qdots for both normal streptavidin conjugates, indicating that they had cross-linked to multiple molecules, even at the 0.2 % biotin concentration shown in Figure 4 - supplement 3. At 1.0 % biotin, there was even a larger population of slow diffusive qdots, i.e., cross-linked qdots. By contrast, in the case of mSA qdot conjugates, we found that most qdot conjugates were freely diffusive at 0.2 % as well as at 1.0 % biotin concentration; that is, mSA-qdots do not show a slow diffusive population of labeled molecules at either 0.2 % or 1.0 % concentrations. These observations showed that normal streptavidin frequently cross-links to multiple molecules while monomeric streptavidin did not.

In order to further examine cross-linking of normal and monomeric streptavidin, we compared the run length of kinesin (K432, biotin at C-terminus). It is well known that when the cargo (in this case the streptavidin-qdot conjugate) binds to multiple motor proteins, the cargo will be transported over a longer distance than when the cargo binds to a single motor protein. (Beeg et al., 2008; Furuta et al., 2013) A single motor protein transports cargo approximately 1 um, while multiple motor proteins can walk farther than 1 um. Thus, if we compare the run length of kinesin using SA-qdots and mSA-qdots at different ratios of kinesin and qdots, then we can examine the possibility that one qdot is binding to multiple proteins versus binding to a single protein. For accuracy of labeling, we want to confirm that binding is to a single protein and a shorter kinesin run length will indicate this.

In the case of mSA-qdots, we found the run length was the same as that of a single kinesin at a ratio of 2:1 (2 kinesin molecules to 1 qdot). As expected (Figure 4 - supplement 4), the run length was not changed even when the ratio of kinesin to qdots was increased to 6:1. This means that mSA-qdots do not bind to multiple motor proteins. By contrast, in the case of SA-qdots, the run length of kinesin was more than the typical run length of kinesin, as shown in figure S11. Even at a ratio of 2:1, the histogram of run lengths shows that some kinesin walked longer than the typical kinesin run length. At 6:1, we found that the run length significantly increased, indicating cross-linking. These results mirror the results we obtained in our first experiment about diffusion rates of labeled molecules in the lipid bilayer.

**Figure 2 - figure supplement 1.**
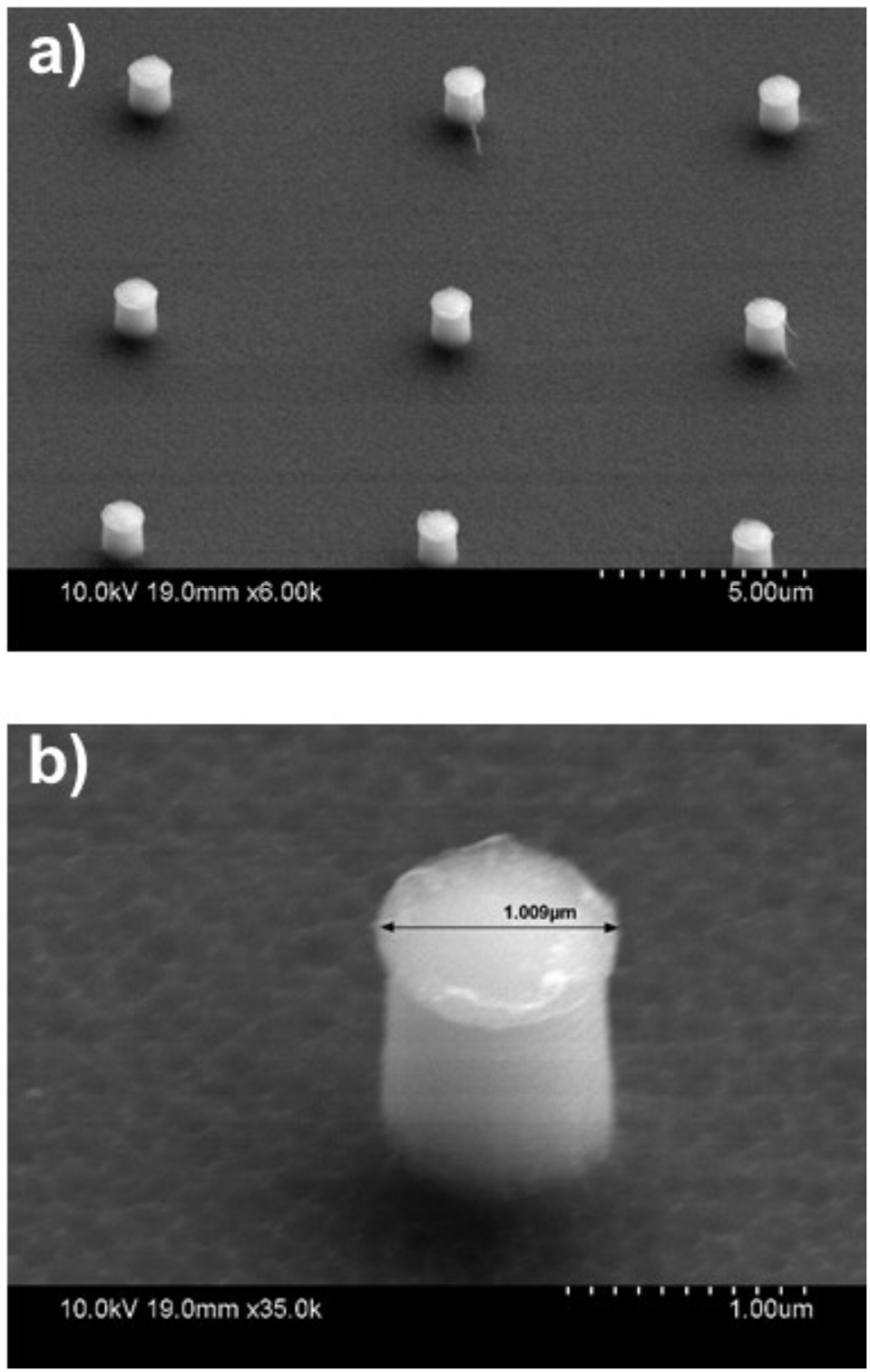
TEM image for Si mask which has fiduciary markers. Fiduciary markers are every 8 μm or 16 μm and total pattern size is 18 mm and 18 mm. Each marker size of the round column is 1 μm height and 1 μm diameter. We then made the mold using PDMS and stamped the pattern on the coverslip.

**Figure 2 - figure supplement 2.**
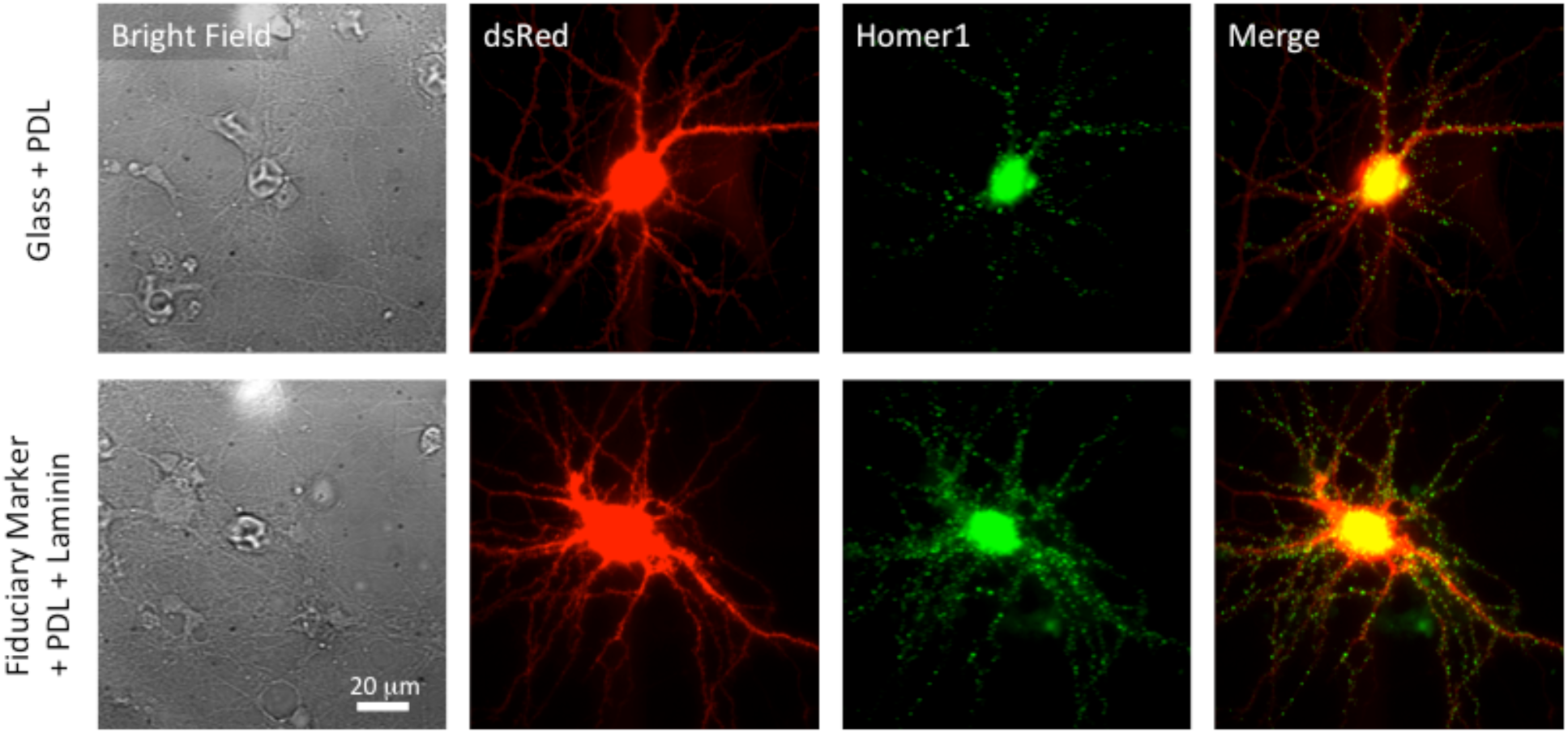
The upper panel represents cultured neurons on the poly-D-lysine (PDL) coated glass coverslip. The first column is bright field images. Red fluorescence is free ds-Red and green is homer1-mGEOS. The last column is overlay of ds-Red and Homer1-mGEOS. The lower panel is cultured neurons on the fiduciary marked coverslip with PDL and laminin coating. Before coating PDL/Laminin, fiduciary marked coverslips were treated with oxygen plasma to make hydrophilic surface. The neuron cultured on the fiduciary marked coverslip looks very similar to the neuron that cultured on the glass coverslip in terms of their shape and number spines. All images are taken with x60 water immersion Nikon objective lens (Plan Apo VC 60x WI).

**Figure 2 - figure supplement 3.**
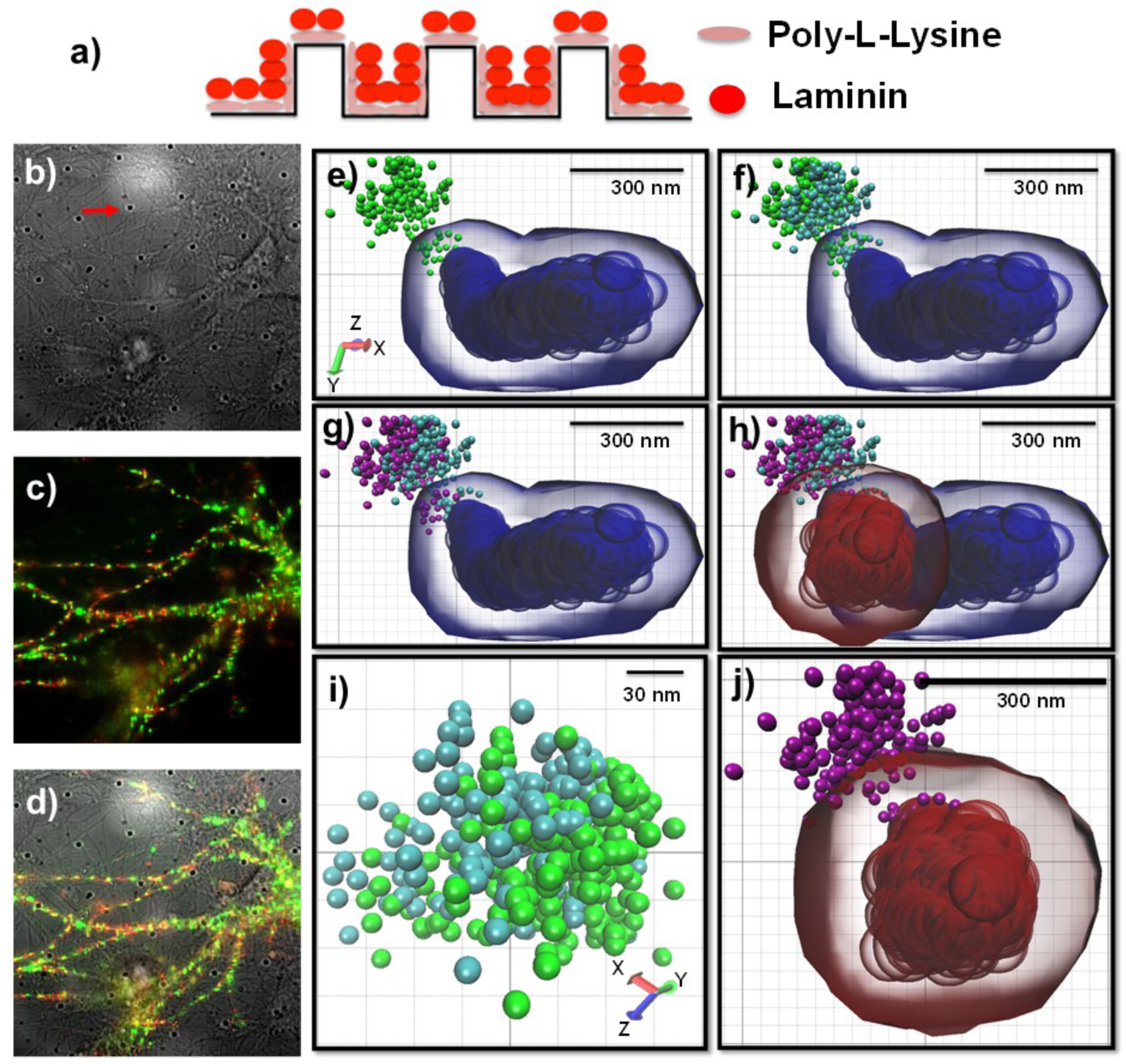
Schematics for culturing neurons on the fiduciary marked coverslips and presentative images for correcting stage drift and chromatic aberration. a) Fiduciary marked coverslips need to be coated with poly-(L/D)-lysine and laminin, sequentially. These coating helps neurons adhere on the surface. b) a bright field image which shows neurons on the fiduciary marked coverslips. Periodic black dots are fiduciary markers as indicated by a red arrow. c) fluorescence images for neurons. There is no auto-fluorescence at the illumination of visible wavelength. d) overlay image of c) and d). e) a representative reconstructed image for a synapse (homer1, blue) and the trace of a receptor (AMPA receptor, green) without correcting stage drift and chromatic aberration. f) corrected chromatic aberration. Using the mapping function to correct chromatic aberration, the red color channel transposed to the green channel. Cyan color balls represented the trace of AMPA receptors after correcting chromatic aberration. g) Purple balls represented the trace of a receptor after correcting stage drift. This trace (purple balls) has minimal error caused by stage drift and chromatic aberration. h) correcting stage drift for the image of a synapse. Blue color shows before correction and red color shows after correcting stage drift. i) zoom-in image showing difference between before and after correcting chromatic aberration.

**Figure 2 - figure supplement 4.**
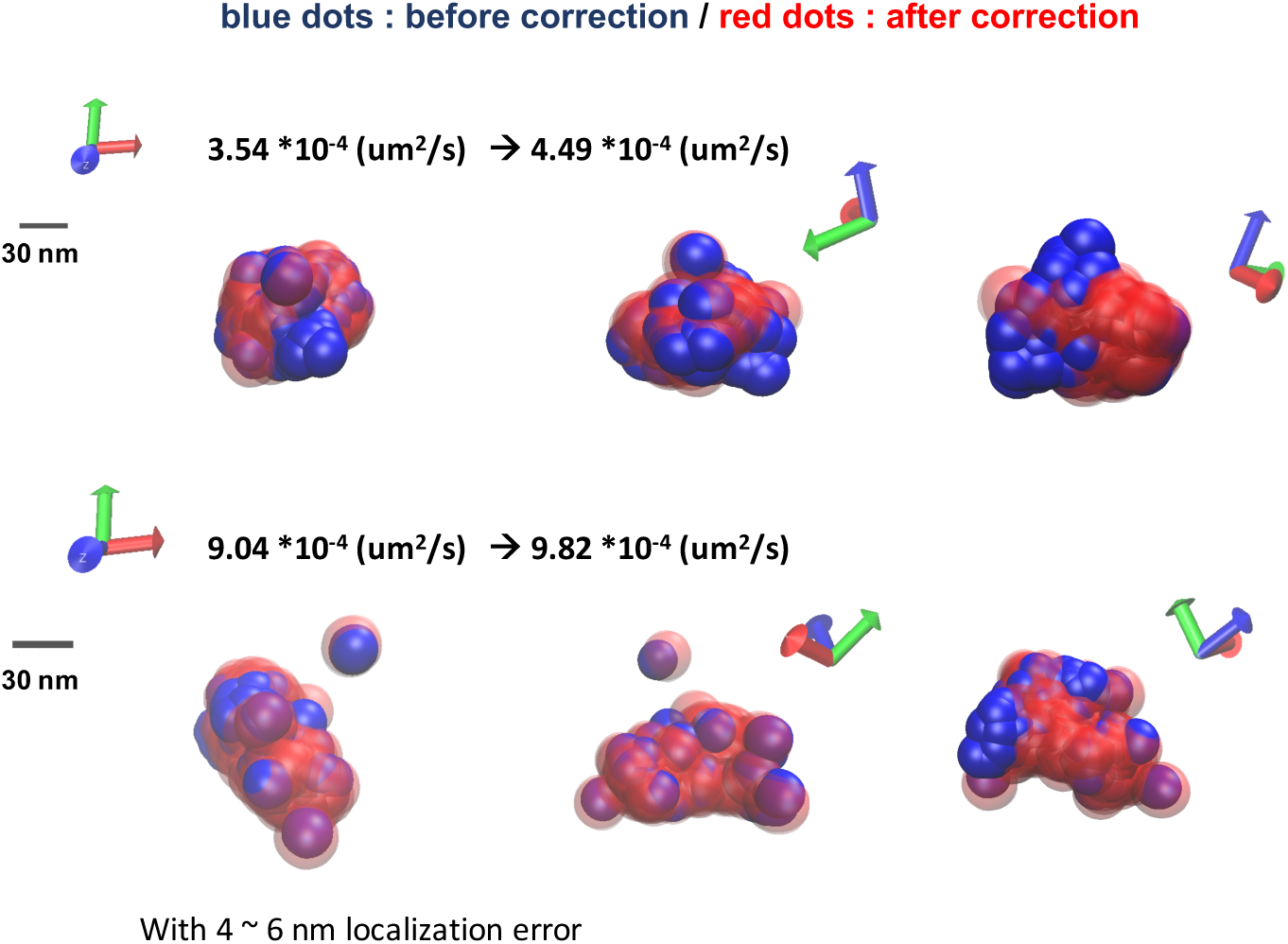
The difference of diffusion coefficient with and without correcting the stage drift. Even though the tracking AMPA receptors takes for 50s, the stage drift affects measuring diffusion coefficient. At maximum, there is more than 10 % difference between before and after correcting the stage drift. With astigmatism for three dimensional imaging, the localization precision is about 4 nm for x and 6 nm for y.

**Figure 2 - figure supplement 5.**
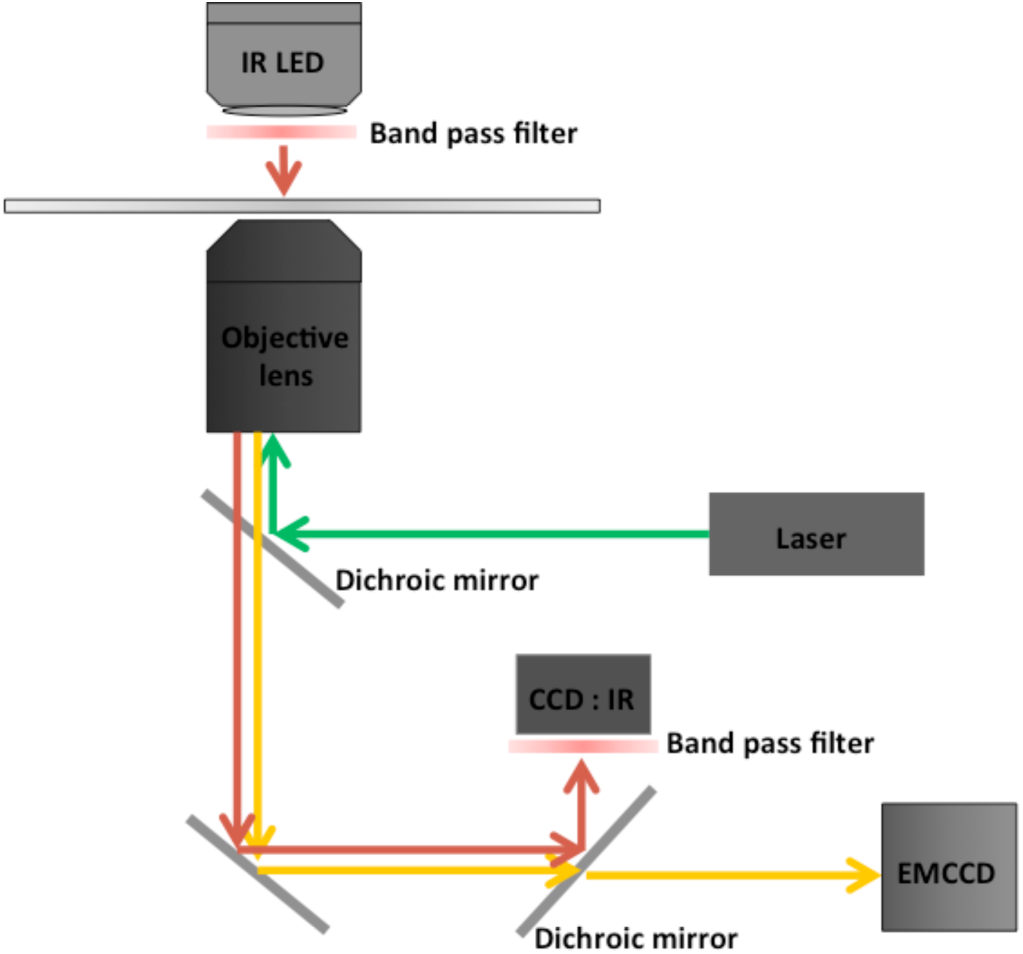
Commercially available microscopes have a perfect focus system (PFS), which normally uses IR LED light, in order to reduce the drift along z-axis. In the case of Nikon (*Ti-eclipse*), the wavelength of the LED is 870 nm with 800 nm short pass dichroic mirror. We then selected 760 nm to avoid interference to the PFS. The band-width of the 760 nm LED (ELJ-750-629, Roithner-LaserTechnik GmbH) is from 725 nm to 775 nm and we added the band pass filter (769/41, OD6 blocking) at LED as well as at the IR camera in order to reduce the interference between the LED for the PFS and the IR LED for fiduciary markers.

**Figure 2 - figure supplement 6.**
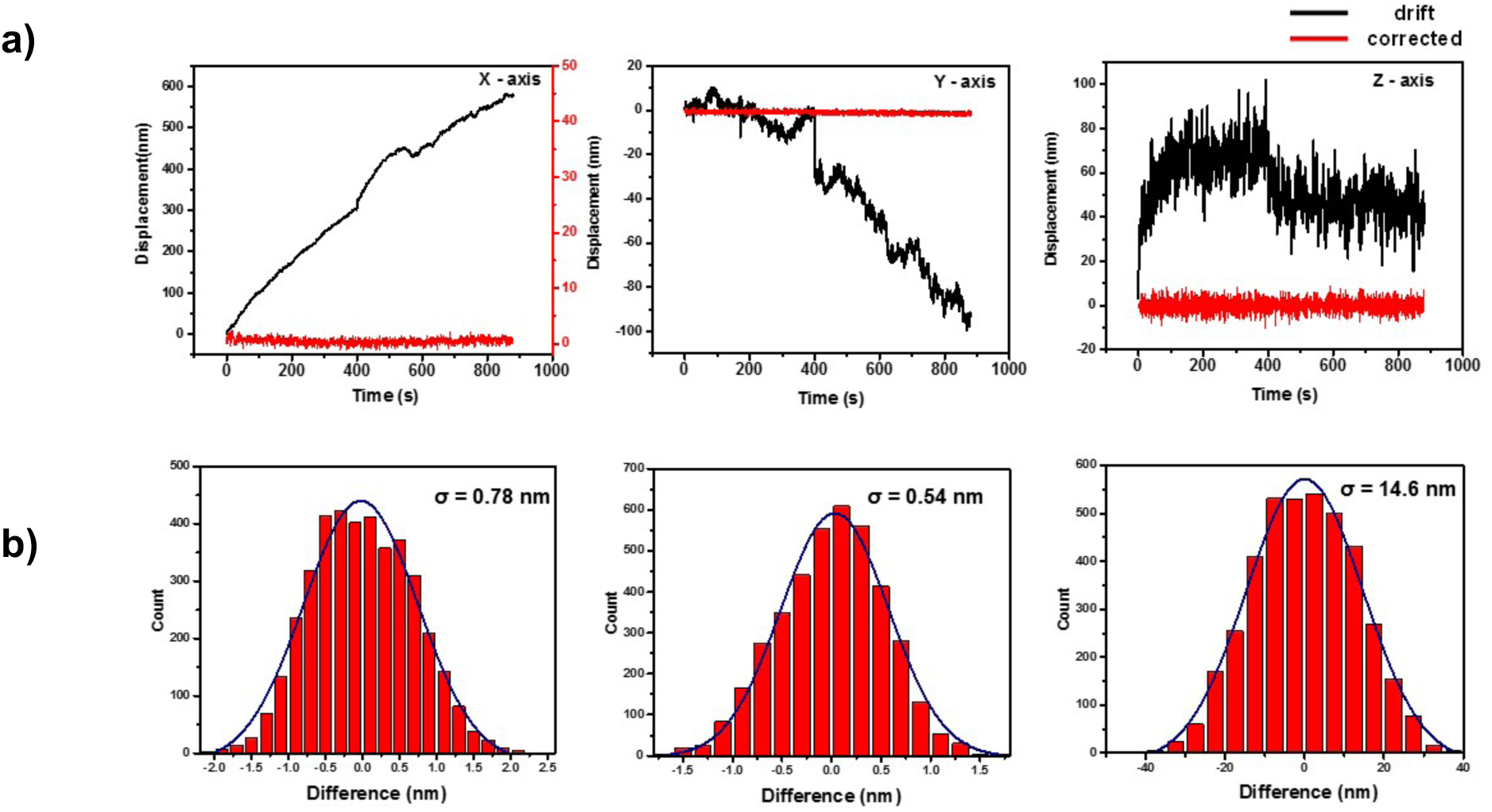
Traces of the stage drift and the precision of the correction. a) All three axis show significant stage drift over time. Black lines represent the stage drift and the stage drift was corrected by using fiduciary markers. Red lines represent the corrected traces of the stage. b) The precisions are less than 1 nm for x and y axis and 14.6 nm for z-axis.

**Figure 2 - figure supplement 7.**
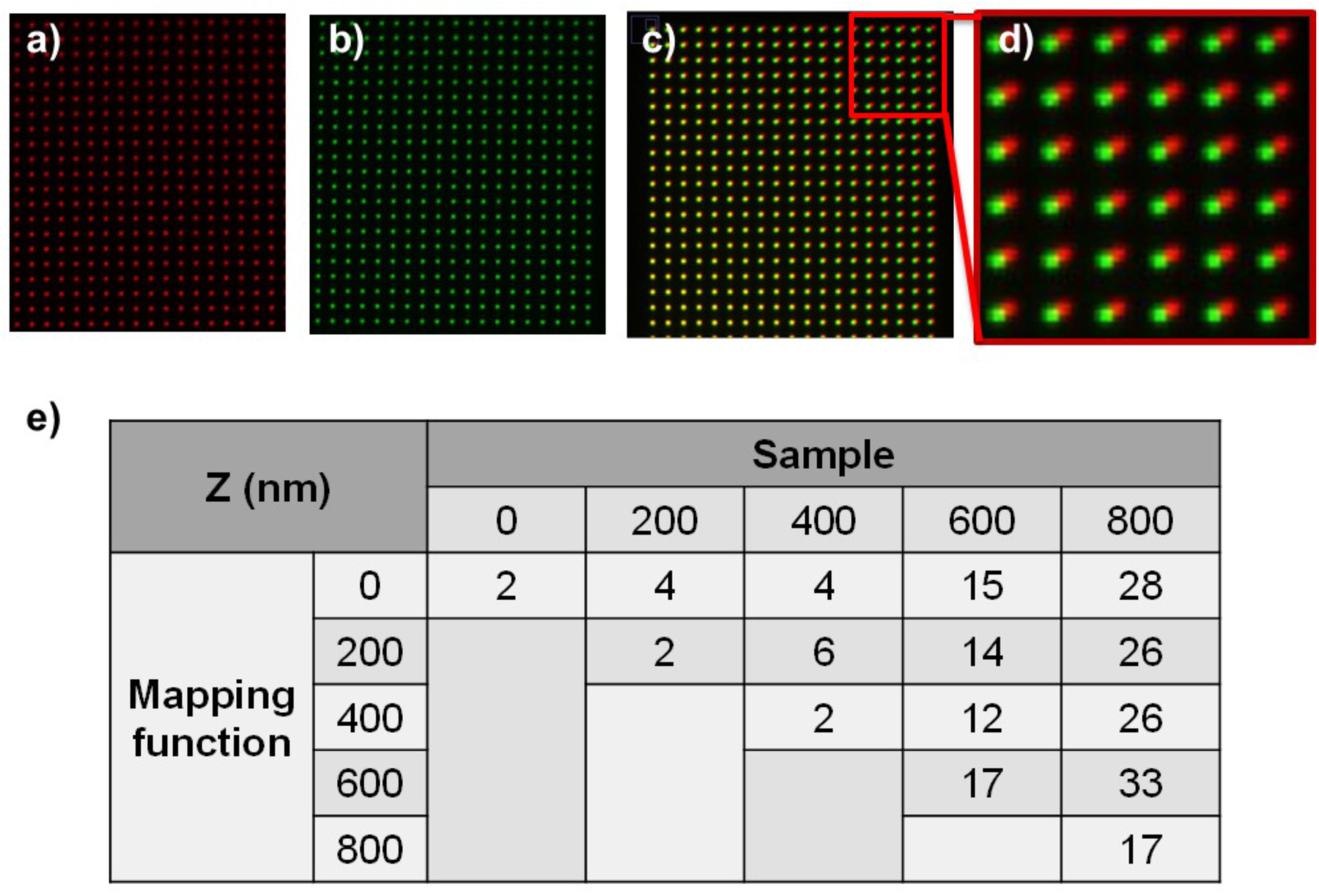
Correcting chromatic aberration using nanohole pattern. We made 100 nm size hole every 1.5 μm on the silver coated glass coverslip. When illuminating white light through the nanohole pattern, each color image can be taken using a band pass filter.

**Figure 4 –figure supplement 1.**
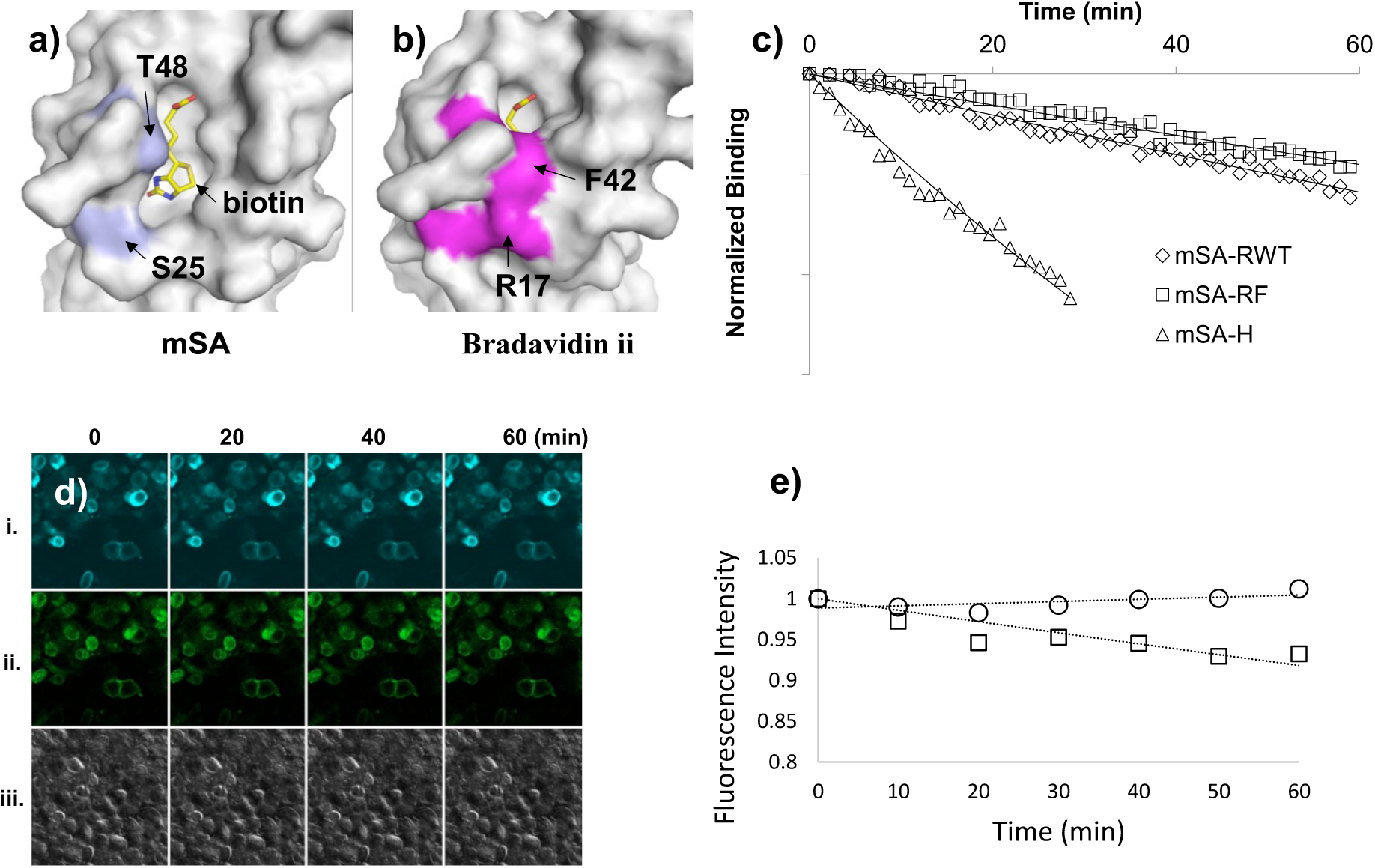
Characterization of monomeric streptavidin and conjugation with small quantum dots. a) The biotin binding pocket of mSA is exposed to the solvent, which results in rapid dissociation of bound biotin. In contrast, b) the binding pocket of bradavidin ii is surrounded by R17 and F42, which surround the biotin and limit access of water molecules. Double mutations, S25R and T48F, in mSA may similarly shield bound biotin from solvent molecules and slow the rate of dissociation. c) The dissociation rate, k_off_, of mSA variants was measured by fluorescence polarization spectroscopy using biotin-fluorescein as the ligand. Fluorescence polarization from the ligand is reduced upon dissociation from mSA. Normalized binding is computed by subtracting the polarization of free dye, *P_free_*, and dividing by the polarization difference between bound and unbound states, (*P_bound_* – *P_free_*). Fitting normalized polarization to an exponentially decaying curve *e^−k_off_*t^* results in the off rate, k_off_. d) To test for cell labeling of mSA, mSA-RF-GFP was used to label biotinylated AP-CFP that transfected on HEK293. Images of i/ii/iii show CFP, GFP, and bright field images. The fluorescence intensity of GFP and CFP does not show much difference. d) and e) Test specificity of mSA-RF-eGFP to AP-CFP on HEK293. After the cells were fixed, mSA-RF-EGFP was added for 30 min to label biotinylated proteins. Unbound mSA-RF—EGFP was washed away and the labeled cells were imaged for i. CFP or ii. GFP fluorescence every 10 min for 1 hr. iii. DIC, differential interference contrast. The total fluorescence intensity of each image was computed at different time points by importing RGB jpg images into MatLab and by summing over green and blue channels (CFP) or over green channel only (GFP).

**Figure 4 –figure supplement 2.**
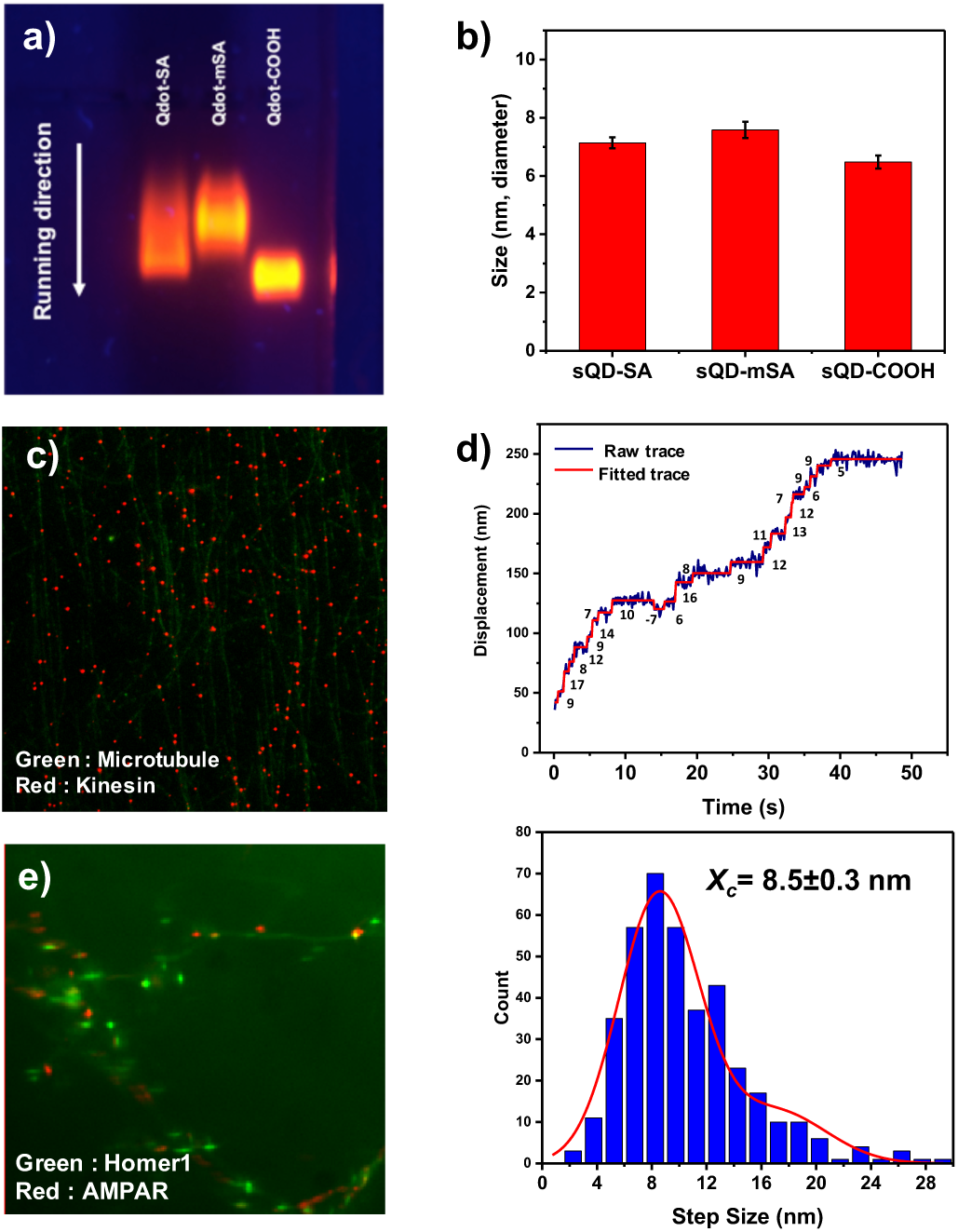
a) Agarose gel image for SA-, mSA-, and carboxylated- small qdots and b) histogram for their size. c) Motor proteins (kinesin) labeled by mSA-small qdot (red dots) are on microtubules (green, HiLyte 488). d) Kinesin walking trace and their step size which shows 8.5 nm. e) mSA-small qdots label AMPA receptors on neurons (green fluorescence is Homer1). mSA-small qdots are specifically bound to AMPA receptors (red).

**Figure 4 –figure supplement 3.**
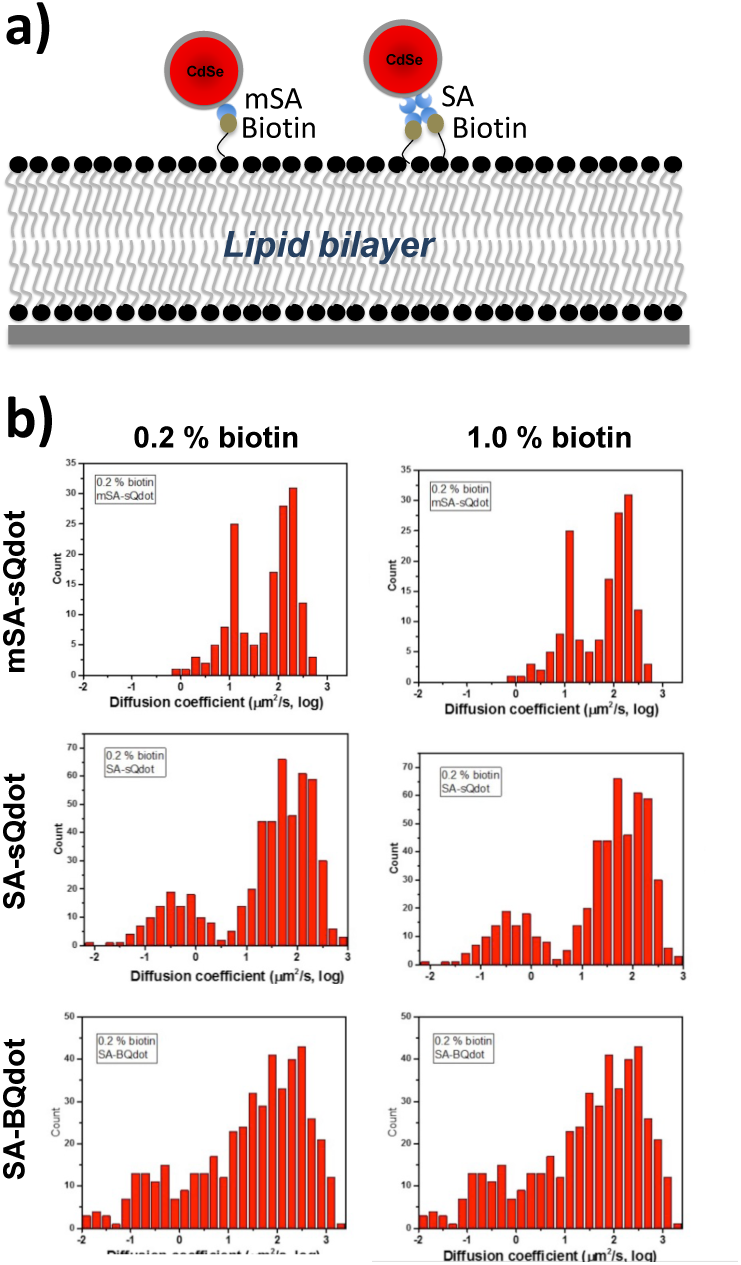
Measuring the diffusion coefficient of labeled qdots on the lipid bilayer in order to examine the possibility of cross-linking. a) represents the schematics of cross-linking of streptavidin(SA) conjugated qdots. SA conjugated qdots have more chance to cross-link multiple target molecules on a single qdots because a SA has four binding sites, but a monomeric SA has only one binding site. Thus, monomeric streptavidin intrinsically excludes binding multiple molecules on a single qdot. b) histograms of diffusion coefficients measured at different conditions which are at 0.2 % and 1.0 % biotin concentration of lipid bilayer using three different qdots, such as SA-Bqdot(commercial qdots), SA-small qdots, and monomeric SA-small qdots.

**Figure 4 –figure supplement 4.**
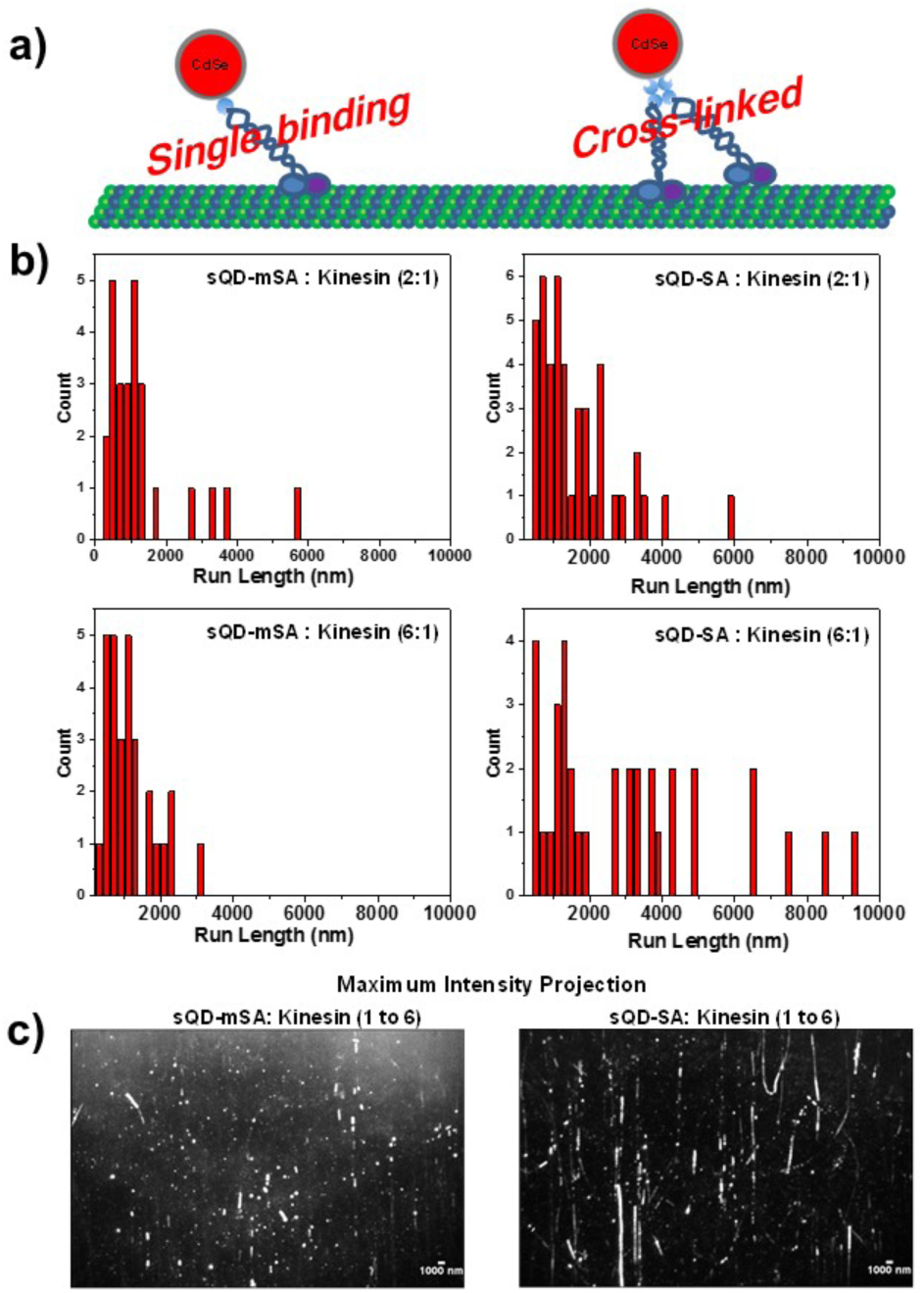
Measurement the run-length of kinesin using mSA-small qdot and SA-small qdot. a) schematics to show the possibility of crosslinking multiple kinesin on qdots. Since a mSA is monovalent, there is no possibility to bind multiple kinesin. By contrary, a streptavidin has four binding sites for biotin, so that, even though each small qdot has only one streptavidin, SA-small qdots can bind more than one kinesin. b) the measurement of the run length of kinesin at two different concentration ratio between qdots and kinesin. c) kymograph for kinesin walking traces with 1:6 (small qdot: kinesin) ratio.

**Figure5 –figure supplement 1.**
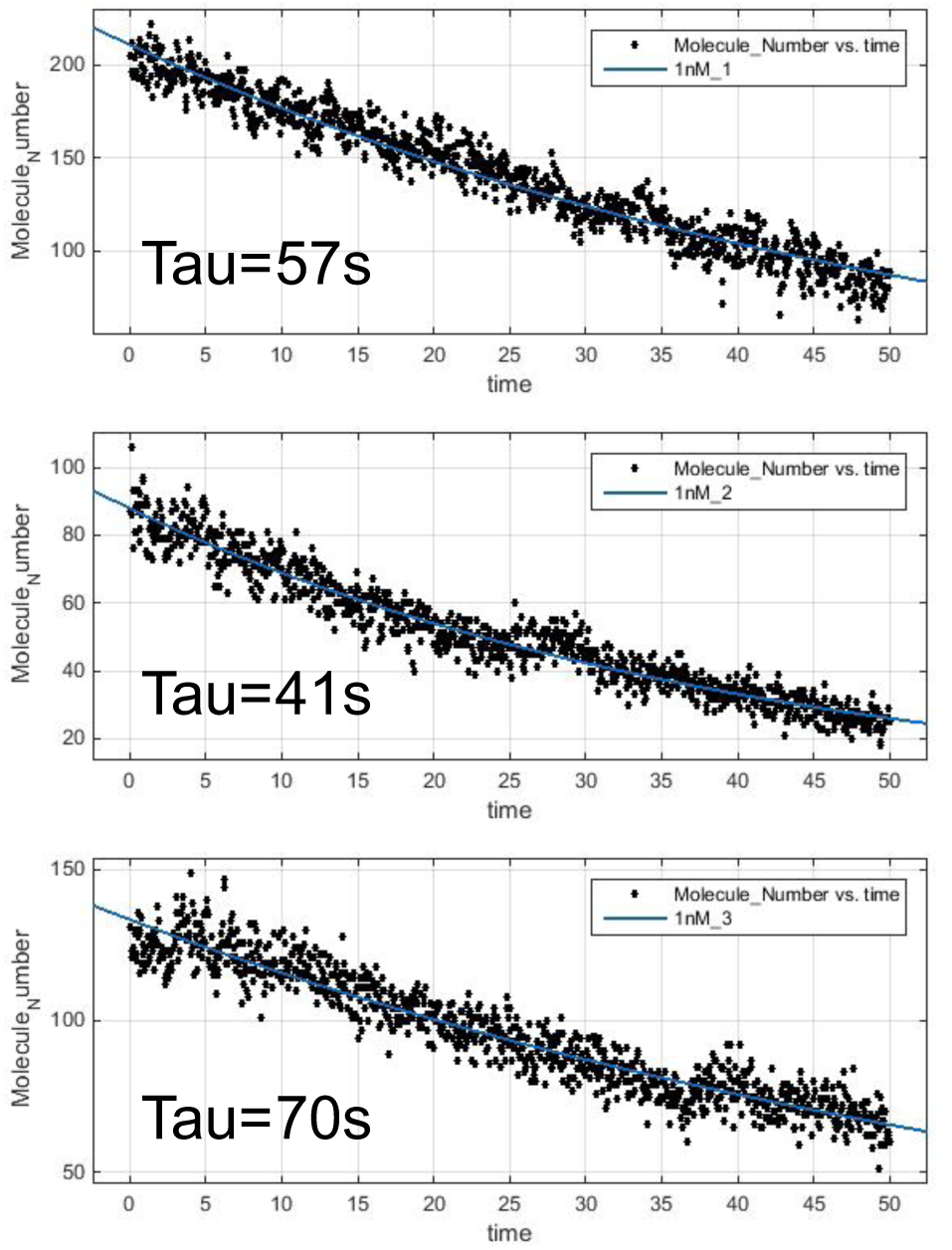
Measurement of the photobleaching time of SA-atto647N. At 1 nM of SA-atto647N, the average photobleaching time is about 61 second. Thus, we were able to measure the diffusion coefficient of AMPA receptors using SA-atto647N.

**Figure 5 –figure supplement 2.**
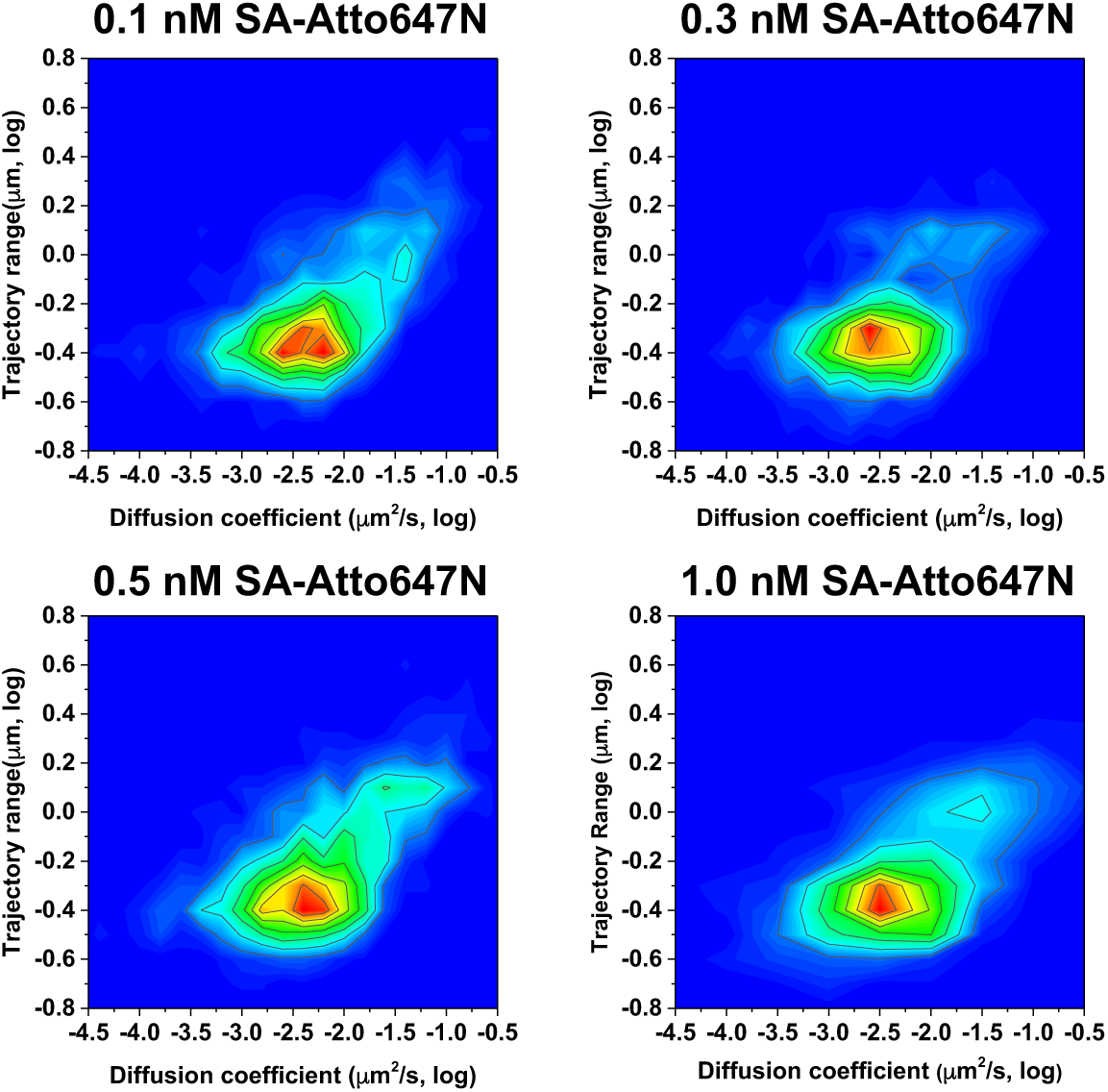
Two dimensional heat maps for the diffusion coefficient and trajectory range measured at various concentrations (0.1, 0.3, 0.5, and 1.0 nM) of SA-Atto647N. The distribution of diffusion coefficient and trajectory range does not change at different concentrations.

**Figure5 –figure supplement 3.**
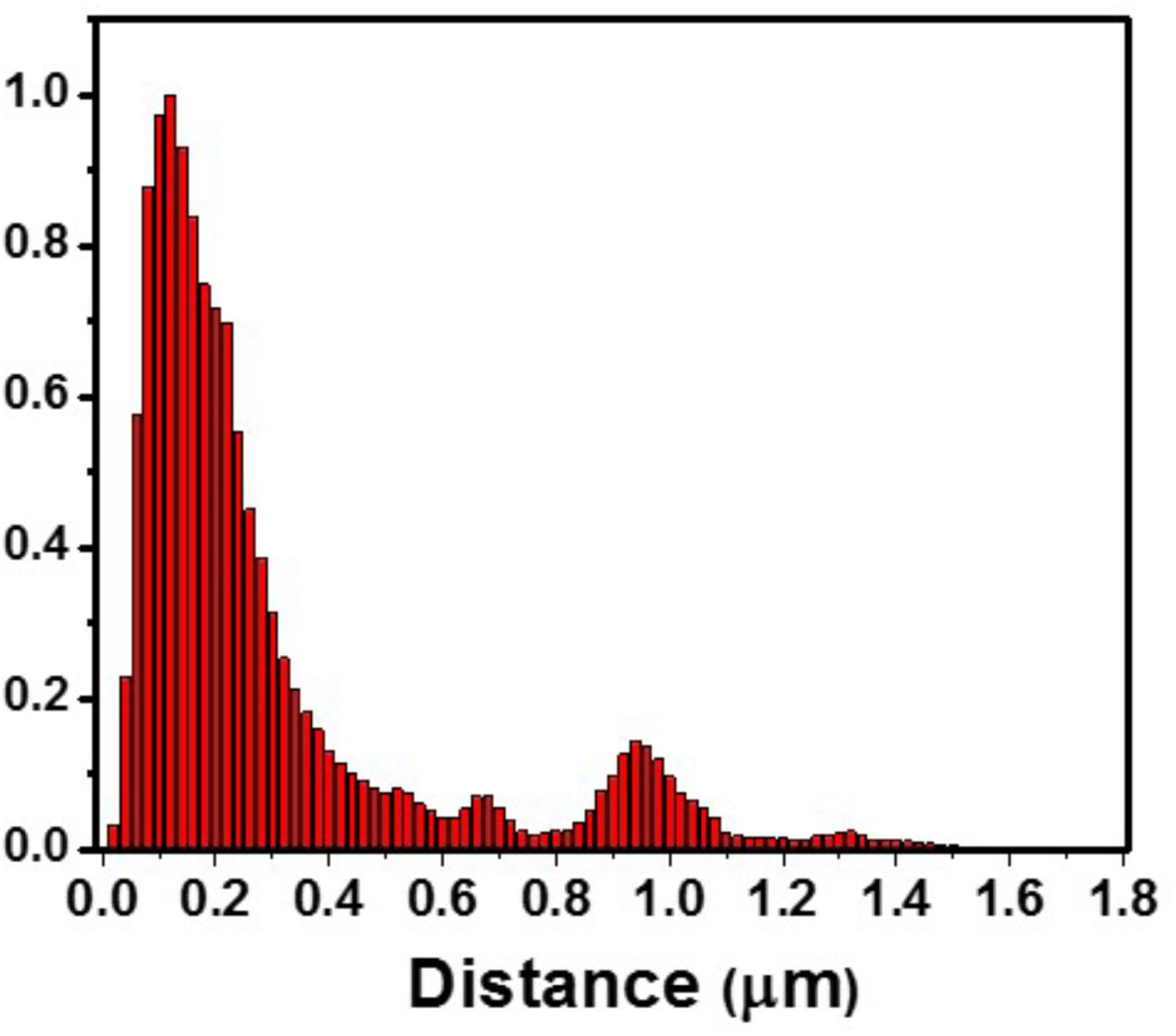
Measured the distance between the receptors and the center of Homer1c. For organic dyes, 85% of receptors are localized within 0.5 μm.

